# Shared transcriptional control and disparate gain and loss of aphid parasitism genes and loci acquired via horizontal gene transfer

**DOI:** 10.1101/246801

**Authors:** Peter Thorpe, Carmen M. Escudero-Martinez, Peter J. A. Cock, D. Laetsch, Sebastian Eves-van den Akker, Jorunn I.B. Bos

**Affiliations:** Cell and Molecular Sciences, The James Hutton Institute, Invergowrie, Dundee, DD2 5DA, UK; Information and Computational Sciences, The James Hutton Institute, Invergowrie, Dundee, DD2 5DA, UK; Dundee Effector Consortium, The James Hutton Institute, Invergowrie, Dundee, DD2 5DA, UK; Institute of Evolutionary Biology, University of Edinburgh, EH9 3FL, UK; Division of Plant Sciences, School of Life Sciences, University of Dundee, Dundee; Biological Chemistry, John Innes Centre, Norwich Research Park, Norwich NR4 7UH

**Keywords:** aphids, effectors, genome evolution, shared transcriptional control, horizontal gene transfer

## Abstract

**Background:** Aphids are a diverse group of taxa that contain hundreds of agronomically important species, which vary in their host range and pathogenicity. However, the genome evolution underlying agriculturally important aphid traits is not well understood.

**Results:** We generated highly-contiguous draft genome assemblies for two aphid species: the narrow host range *Myzus cerasi*, and the cereal specialist *Rhopalosiphum padi*. Using a *de novo* gene prediction pipeline on both these genome assemblies, and those of three related species (*Acyrthosiphon pisum, D. noxia and M. persicae*), we show that aphid genomes consistently encode similar gene numbers, and in the case of *A. pisum*, fewer and larger genes than previously reported. We compare gene content, gene duplication, synteny, horizontal gene transfer events, and putative effector repertoires between these five species to understand the genome evolution of globally important plant parasites.

Aphid genomes show signs of relatively distant gene duplication, and substantial, relatively recent, gene birth, and are characterized by disparate gain and loss of genes acquired by horizontal gene transfer (HGT). Such HGT events account for approximately 1% of loci, and contribute to the protein-coding content of aphid species analysed. Putative effector repertoires, originating from duplicated loci, putative HGT events and other loci, have an unusual genomic organisation and evolutionary history. We identify a highly conserved effector-pair that is tightly genetically-linked in all aphid species. In *R. padi*, this effector pair is tightly transcriptionally-linked, and shares a transcriptional control mechanism with a subset of approximately 50 other putative effectors distributed across the genome.

**Conclusions:** This study extends our current knowledge on the evolution of aphid genomes and reveals evidence for a shared control mechanism, which underlies effector expression, and ultimately plant parasitism.

## Background

Among the over 5000 aphid species described to date, about 250 are important agricultural pests [1]. These aphid species are highly diverse with regards to many phenotypic and ecological traits. Interestingly, whilst host specialization on a single or few plant species is common, some aphid species have evolved to infest a wide range of plant species, including from different families. How interactions with biotic factors have shaped aphid diversity is a complex and unanswered question. With increasing numbers of aphid genomes becoming available, it is possible to interrogate the evolution of genes that are predicted to play a role in aphid-environment interactions, such as host parasitism.

Genome sequences have become available for four different aphid species, *Acyrthosiphum pisum* (pea aphid) [2], *Myzus persicae* (green-peach aphid) [3], *Diuraphis noxia* (Russian wheat aphid) [4], and most recently, *Aphis glycines* (soybean aphid) [5]. Already, this has led to important discoveries, such as the association of duplicated gene cluster transcriptional plasticity in the broad host range *M. persicae* with colonization of diverse host species [3], and the discovery that genes involved in carotenoid biosynthesis in the pea aphid were acquired by horizontal gene transfer (HGT) from fungi [6]. Screening of the pea aphid genome for genes acquired by HGT from bacteria identified only 12 candidates, of which at least 8 appeared to be functional based on expression data [7]. HGT could therefore have played a role in the acquisition of novel important aphid traits, but the extent of its impact on aphid genome evolution, and host-parasite interactions, remains unclear.

Recent progress in the field revealed that a molecular dialogue takes place between plants and aphids leading to activation of plant defences in resistant plants (reviewed by [8]), or the suppression of host defences and/release of nutrients in susceptible plants [9] [10] [11]. Aphid effectors, which are molecules delivered inside host plant cells and the apoplast during probing and feeding, play an important role in the infestation process in that they contribute to host susceptibility by targeting host cell processes (reviewed in [12] and [13]) [14]. Recent progress in aphid transcriptomics and proteomics facilitated the identification of effectors in several important species [15] [16] [17] [18] [19], and revealed overlap and diversity between species [20]. Expanding comparative analyses of aphid effectors to the genome level promises to provide new insight into their evolution. For example, in the case of plant parasitic nematodes and filamentous plant pathogens, effectors tend to be located in gene-sparse regions, that are repeat-rich to allow for adaptive evolution [21] [22].

In this study, we sequenced the genomes of *Myzus cerasi* (black-cherry aphid), which is closely related to *M. persicae* but in contrast has a limited host range, and *Rhopalosiphum padi* (bird-cherry oat aphid), which is a cereal specialist. Together with three previously published aphid genomes (*A. pisum, D. noxia* and *M. persicae*), we compare gene content, duplication, HGT events, and effector repertoires. Importantly, our gene model (re-)prediction approach revealed that the different aphid genomes have more consistent gene number than previously reported [3] [2], between 25,726 and 28,688 genes predicted across the different genomes. A combination of gene duplication, gene birth, as well as HGT events has shaped aphid genomes, and contributed to the acquisition of predicted aphid effector genes. Strikingly, we found that expression of a subset of these aphid effector genes is tightly co-regulated, reflecting the presence of an unknown transcriptional control mechanism that likely underpins plant- parasitism.

## Results and Discussion

We sequenced the genomes of a clonal line of *M. cerasi* established on secondary host species *Barbarea verna* (Land Cress) and of a clonal line of *R. padi* established on *Hordeum vulgare* (Barley) using Illumina 2X250 bp pair-end libraries (and 2X150bp for *M. cerasi*) to a depth of 233x and 129x, respectively. Using these data, the genome of *M. cerasi* was assembled to 406 Mbp contained in 49,349 contigs and the *R. padi* genome assembled to 319 Mbp contained in 15,616 contigs. These assemblies are of a similar size to those reported for *Diuraphis noxia* (393 Mbp; [4]), *Myzus persicae* (347/356 Mbp; [3]), and *Acyrthosiphum pisum* (464 Mbp; [2]) (Table 1). When compared to previously published assemblies of aphid genomes, the *M. cerasi* and *R. padi* genomes are highly contiguous (contig N50 of 19,701 bp and 98,943, respectively) and highly unambiguous (only 54,488 and 194,118 Ns respectively, compared to ~11,500,000, ~41,000,000, and ~98,000,000 Ns for *M. persicae*, *A. pisum* and *D. noxia*, respectively). The endosymbiont genomes (*Buchnera aphidicola*) of *M. cerasi* and *R. padi* were assembled as single contigs of 641,811 bp and 643,950 bp, respectively. Benchmarking Universal Single-Copy Orthologs (BUSCO) [23] and Core Eukaryotic Genes Mapping Approach (CEGMA) [24] were used to estimate an assembly completeness of 80% and 86%, respectively, for *M. cerasi*, and 82% and 93%, respectively, for *R. padi* nuclear genomes (Table 1). The GC content of the *M. cerasi* and *R. padi* genomes (29.9% and 27.8% respectively) is consistent with other aphid genomes [5] [2] [3], and they contain a high proportion of repeat rich and/or transposon-like sequence (Table 1 and Additional file 1: Figure S1). Altogether, these data indicate that, especially in the case of *R. padi*, high quality draft genome assemblies were generated. With a number of aphid genomes available, we are able to perform detailed comparative analyses to understand the evolution of aphid parasitism genes.

**Table 1.**
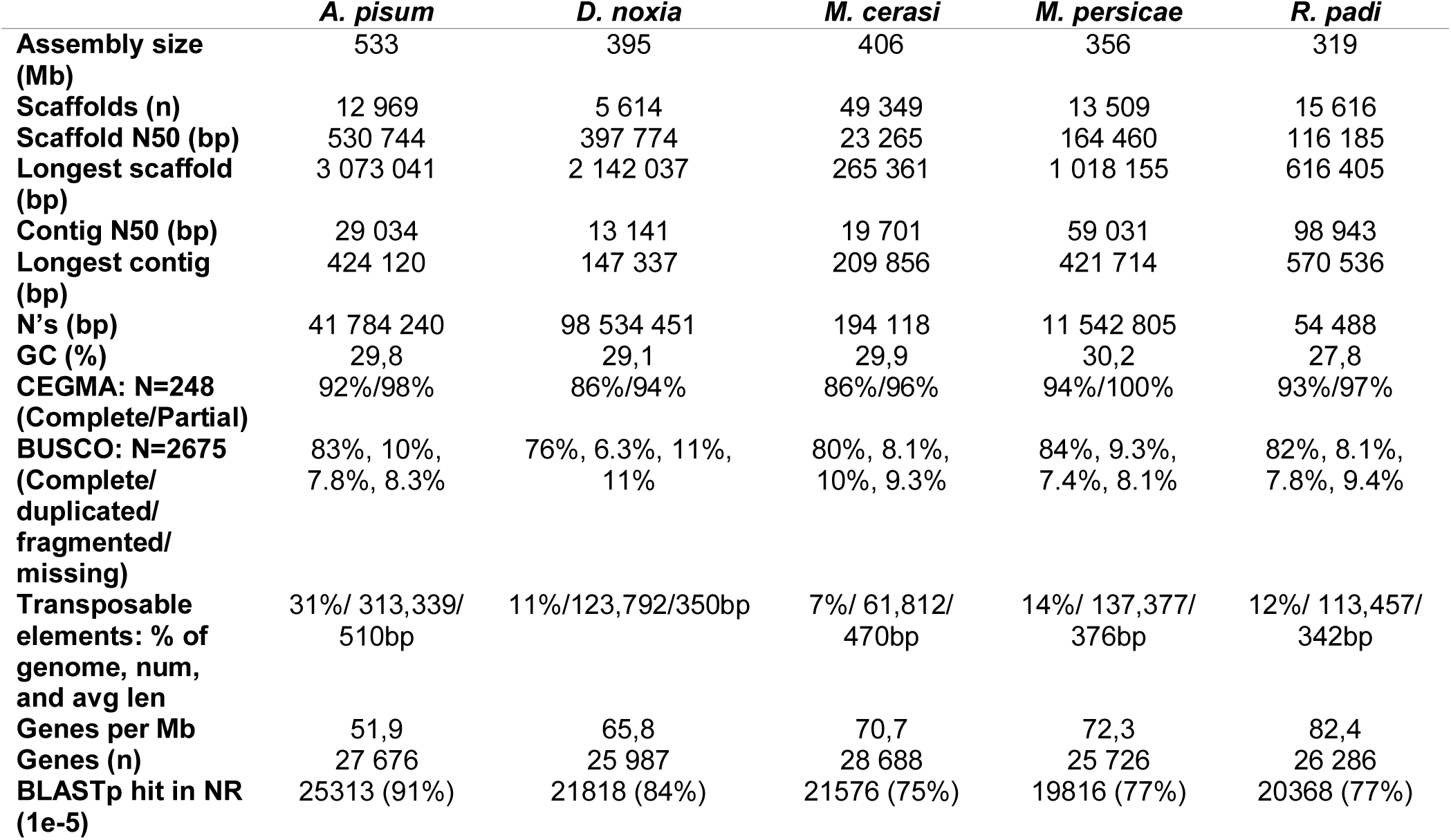
Genome statistics.

|  | *A. pisum* | *D. noxia* | *M. cerasi* | *M. persicae* | *R. padi* |
| --- | --- | --- | --- | --- | --- |
| Assembly size (Mb) | 533 | 395 | 406 | 356 | 319 |
| Scaffolds (n) | 12 969 | 5 614 | 49 349 | 13 509 | 15 616 |
| Scaffold N50 (bp) | 530 744 | 397 774 | 23 265 | 164 460 | 116 185 |
| Longest scaffold (bp) | 3 073 041 | 2 142 037 | 265 361 | 1 018 155 | 616 405 |
| Contig N50 (bp) | 29 034 | 13 141 | 19 701 | 59 031 | 98 943 |
| Longest contig (bp) | 424 120 | 147 337 | 209 856 | 421 714 | 570 536 |
| N's (bp) | 41 784 240 | 98 534 451 | 194 118 | 11 542 805 | 54 488 |
| GC (%) | 29,8 | 29,1 | 29,9 | 30,2 | 27,8 |
| CEGMA: N=248 (Complete/Partial) | 92%/98% | 86%/94% | 86%/96% | 94%/100% | 93%/97% |
| BUSCO: N=2675 (Complete/duplicated/fragmented/missing) | 83%, 10%, 7.8%, 8.3% | 76%, 6.3%, 11%, 11% | 80%, 8.1%, 10%, 9.3% | 84%, 9.3%, 7.4%, 8.1% | 82%, 8.1%, 7.8%, 9.4% |
| Transposable elements: % of genome, num, and avg len | 31%/ 313,339/ 510bp | 11%/123,792/350bp | 7%/ 61,812/ 470bp | 14%/ 137,377/ 376bp | 12%/ 113,457/ 342bp |
| Genes per Mb | 51,9 | 65,8 | 70,7 | 72,3 | 82,4 |
| Genes (n) | 27 676 | 25 987 | 28 688 | 25 726 | 26 286 |
| BLASTp hit in NR (1e-5) | 25313 (91%) | 21818 (84%) | 21576 (75%) | 19816 (77%) | 20368 (77%) |

### Gene model prediction and re-prediction indicate that aphid genomes encode similar gene numbers and, in the case of *A. pisum*, fewer and larger genes than previously reported

To annotate the genome assemblies generated, we initially used the 36,939 *A. pisum* gene models [2], the bundled AUGUSTUS *A. pisum* configuration files [2]. Using this approach, we predicted 35,316 genes for *M. cerasi* (Mc_v0.9), comparable to the 36,939 genes predicted for *A. pisum* ([2] (Additional file 2: Table S1). However, these gene models only describe a minority of the expressed genes, with only 29% of the RNAseq read pairs mapped to the predicted Mc_v0.9 exome. To address this, a subsequent RNAseq-guided *de novo* approach was adopted using BRAKER, generating 28,688 loci for *M. cerasi* (Mc_v1.0). When compared to v0.9 gene models, the *de novo* v1.0 gene models are longer (means of 772 vs 952, respectively), encode almost exactly the same total exome size (27,332,397 vs 27,278,139 nt), contain approximately 35% fewer genes without RNAseq support (8301 vs 5494), and describe more than twice as much of the total RNAseq reads (29% vs 60%, Figure 1A). Comparing the two sets of gene models to one another revealed a markedly different size distribution (Figure 1B). Version 0.9 encodes ~8000 more very short gene models in the size range 0-300 bp than v1.0, and contains 10,411 “unique” loci with no overlap in genomic coordinates with any locus in v1.0 (approximately 29%). The loci unique to v0.9 contribute a larger proportion of the small 0-300 bp gene models than any other gene size category. In contrast to this, the loci unique to v1.0 are evenly distributed across individual size categories and each category is similar to the total proportion in v1.0 (Figure 1B).

**Figure 1.**
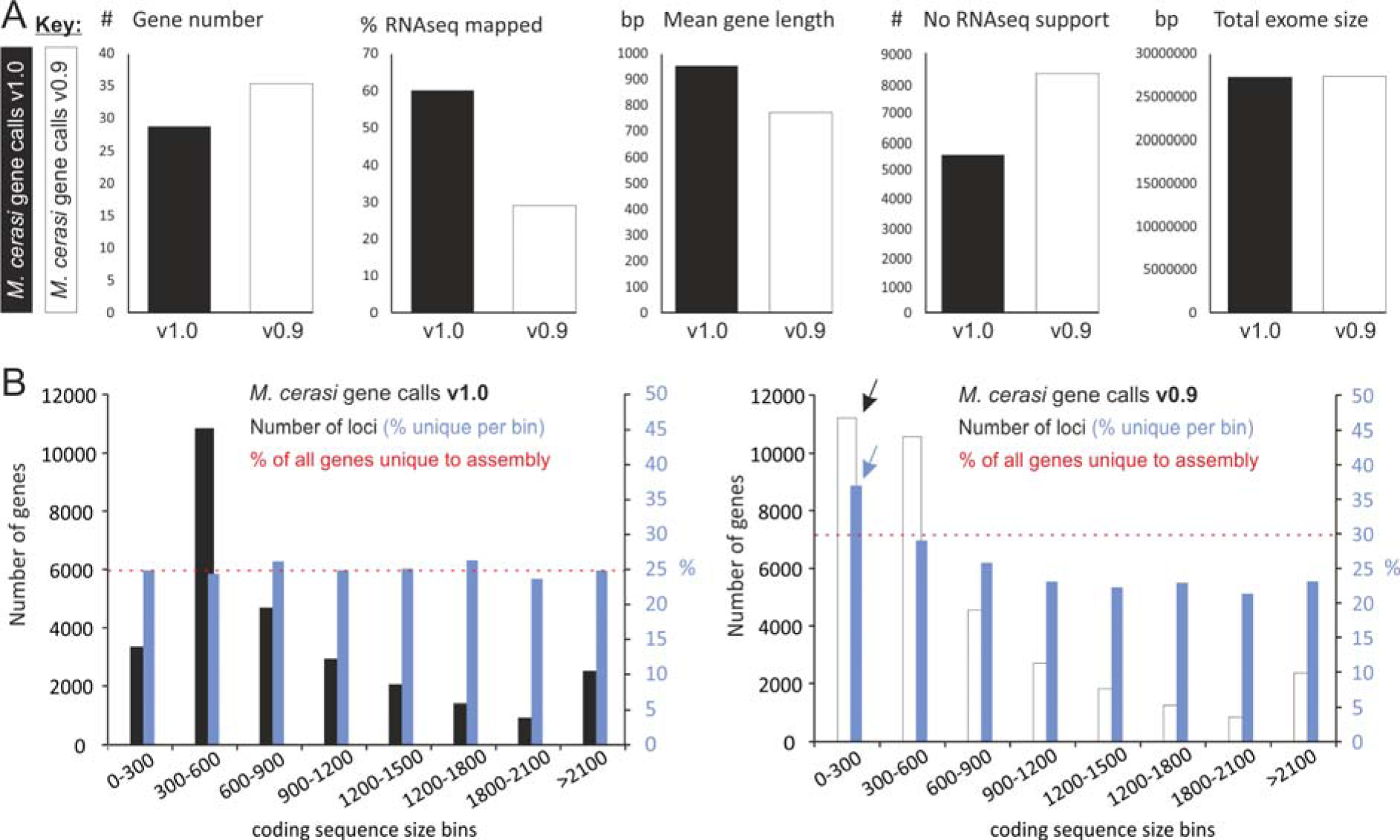
Comparison of *M. cerasi* gene models v0.9 and v1.0. An initial homology- and RNAseq-guided gene model prediction (v0.9, white bars) compared to a subsequent RNAseq-guided *de novo* approach using BRAKER (v1.0, black bars). **A)** v1.0 predictions contained fewer loci, improved mapping of RNAseq reads, were longer on average (mean), had fewer loci with no RNAseq support and yet had an almost identical total exome size to v0.9. **B)** Markedly different frequency distribution of coding sequence length of v1.0 predictions (black) compared to v0.9 predictions (white): v0.9 contains ~8000 very short gene models in the size range 0-300 bp (black arrow). Genes predicted in v1.0 with no corresponding prediction in v0.9 (blue) are evenly distributed across coding sequence size bins. Genes predicted in v0.9 with no corresponding prediction in v1.0 are preferentially contained within the 0-300 bp coding sequence size bin (blue arrow).

Taken together, our results suggest that using gene models of other aphid species to facilitate the annotation of new genomes produces a similar number of loci. However, the majority of these loci are not supported by RNAseq data (even though the RNAseq data were used to facilitate prediction). To avoid propagating errors, we annotated the genome of *R. padi*, and re-annotated all other available aphid genomes, using the RNAseq-guided *de novo* approach described above (Additional file 2: Table S1). Despite being entirely independent, *de novo* annotation produced remarkably consistent gene counts for all aphid species (between 25,726 and 28,688), approximately 25 % less loci overall, and a more complete representation of their individual transcriptomes (Additional file 2: Table S1). Importantly, our approach reduced the possibility that direct comparison of gene content between species is confounded by an inherent bias in different gene prediction methods. Gene models for all species have been made publically available via AphidBase (http://bipaa.genouest.org/is/aphidbase/) and Mealybug (http://mealybug.org/index.html). For the remainder of the manuscript, all comparisons are between the genomes and re/predicted gene content of *M. cerasi, M. persicae, A. pisum, D. noxia*, and *R. padi*.

### Aphid genomes show signs of extensive gene duplication, and recent gene birth

Gene content of aphid genomes is extensively duplicated, and ranges from around 55% multi-copy loci in *M. persicae* to nearly 70% in *A. pisum* (Figure 2A, Additional file 3: Table S2). Although most duplicated loci were classed as “dispersed” rather than “tandem” or “segmental”, we appreciate actual values may slightly deviate from those reported here due to the limit of current assembly contiguity.

**Figure 2.**
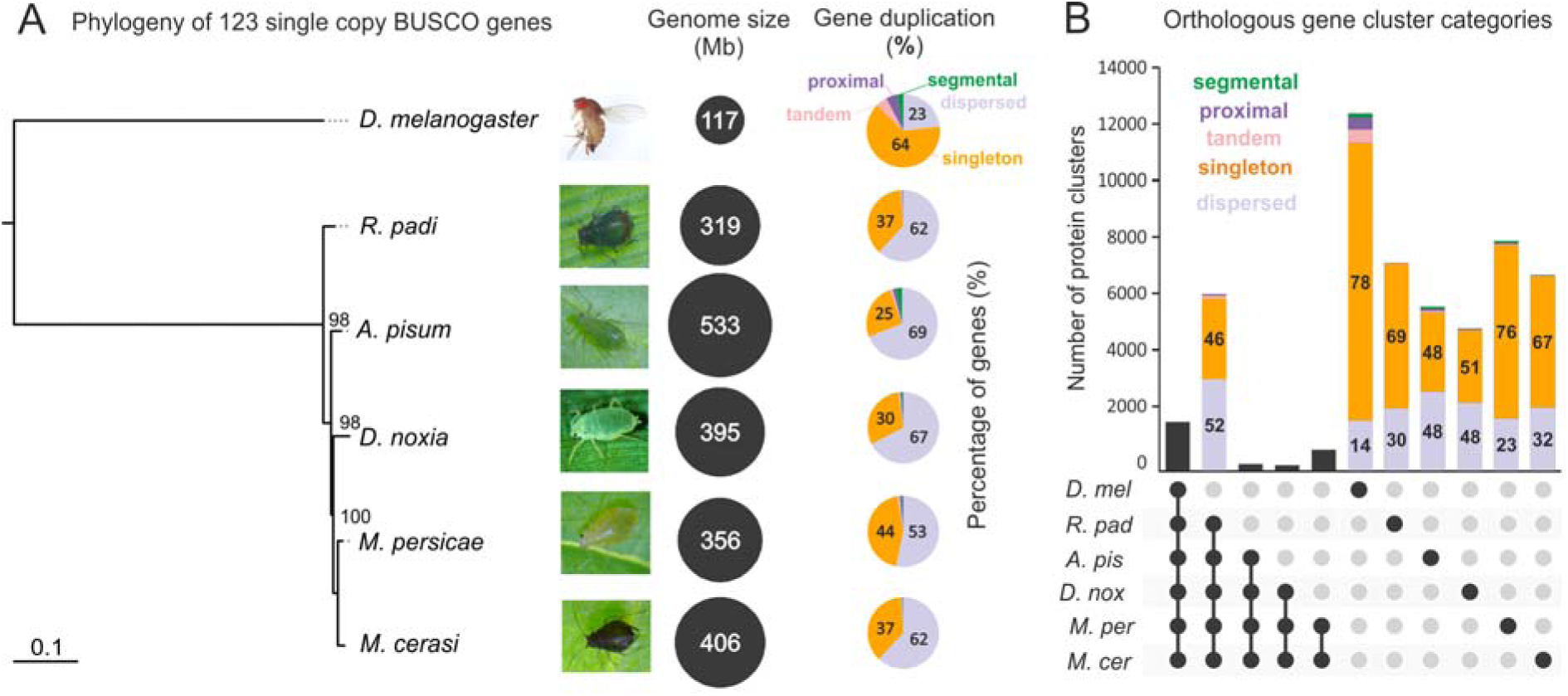
An overview of aphid genomes and gene content. **A**) A multi-gene phylogeny derived from a concatenated alignment of 123 highly-conserved BUSCO nuclear genes classified as single copy in five aphid species (*R. padi,D. noxia, A. pisum, M. persicae*, and *M. cerasi*) and the outgroup model insect *D. melanogaster*. Node values indicate boot strap support of 1000 iterations. For each species, black circles are scaled by genome assembly size, and pie charts are divided by the proportion of gene duplication categories (singleton, dispersed, segmental, proximal, and tandem). **B**) The predicted protein sets of the five aphids were compared to that of the model insect *D. melanogaster*. The histogram shows the number of clusters shared uniquely between the species highlighted below. Selected histograms are divided by the proportion of gene duplication categories, where internal numbers refer to the percentage of that category in that cluster. A total of 6121 clusters contain at least one sequence from each aphid but do not contain any sequence from *D. melanogaster*. Of the genes within these clusters, 54% are duplicated (52% dispersed, 1% tandem and 1% proximal) while 46% are singletons.

To explore the origins of this gene duplication, a robust phylogenetic framework was generated using a multigene phylogeny of 123 highly conserved BUSCO genes present as single copy loci in all aphid genomes tested, and the distant outgroup *Drosophila melanogaster* (Figure 2A). The entire predicted proteomes of all species were clustered based on sequence similarity using MCL, and cross-referenced with the phylogenetic, and gene duplication analyses. This revealed that genes present in clusters with at least one representative from all aphid species but excluding *D. melanogaster* (hereafter referred to as aphid-specific) are primarily multi-copy (Figure 2B). These most likely reflect duplication events that occurred before speciation, followed by retention of multiple copies to present date. In stark contrast, genes present in clusters that exclude all other species (hereafter referred to as species-specific, Figure 2B) are often dominated by single copy loci. Given the relatedness of these aphid species (indicated by short branch lengths in Figure 2A), this observation most likely reflects large scale and relatively recent gene birth in most aphid species, after speciation. Taken together, it is likely that a combination of extensive gene multiplication, and relatively recent gene birth, have shaped the evolution of aphid genomes. Juxtaposed to the recent discovery that duplicated genes play a role in parasitism of the broad host range *M. persicae* [3], the implication of large-scale species-specific gene birth in the context of broad and narrow host-range aphids is intriguing.

### Disparate gain and loss of loci acquired via horizontal gene transfer

To determine whether horizontal gene transfer (HGT) events have contributed to the unusual distribution of gene cluster categories, a systematic genome-wide HGT-identification approach was employed. Putative HGT events were predicted by their ratio of sequence similarity to metazoan and non-metazoan sequences (termed the Alien Index [25] [26] [27]). Using a conservative approach, predicted proteins with an Alien Index > 30 and < 70 % identity to non-metazoan sequences were considered putative HGT, while those with more than 70% identity to non-metazoan sequences were considered putative contaminants, and not further interrogated. Using these criteria, we provide an estimate that ~1-2 % of aphid loci are of non-metazoan origin (between 212 (*M. persicae*) and 338 (*D. noxia*), Figure 3A and Additional file 4: Table S3). While a relatively modest contribution, this estimate expands upon previous reports in both absolute number and donor taxa [7] [6]. HGT events were detected from diverse donor taxa (including plantae, fungi, bacteria, viruses, and other non-metazoan eukaryotes), but were primarily similar to sequences from the fungal and bacterial kingdoms (Figure 3A). Importantly, this approach re-identified carotenoid biosynthesis genes previously characterised as acquired via HGT [6]. Metabolic pathway analysis on all five aphid genomes studied highlights some minor inconsistency in this pathway between species (Additional file 5: Figure S2).

**Figure 3.**
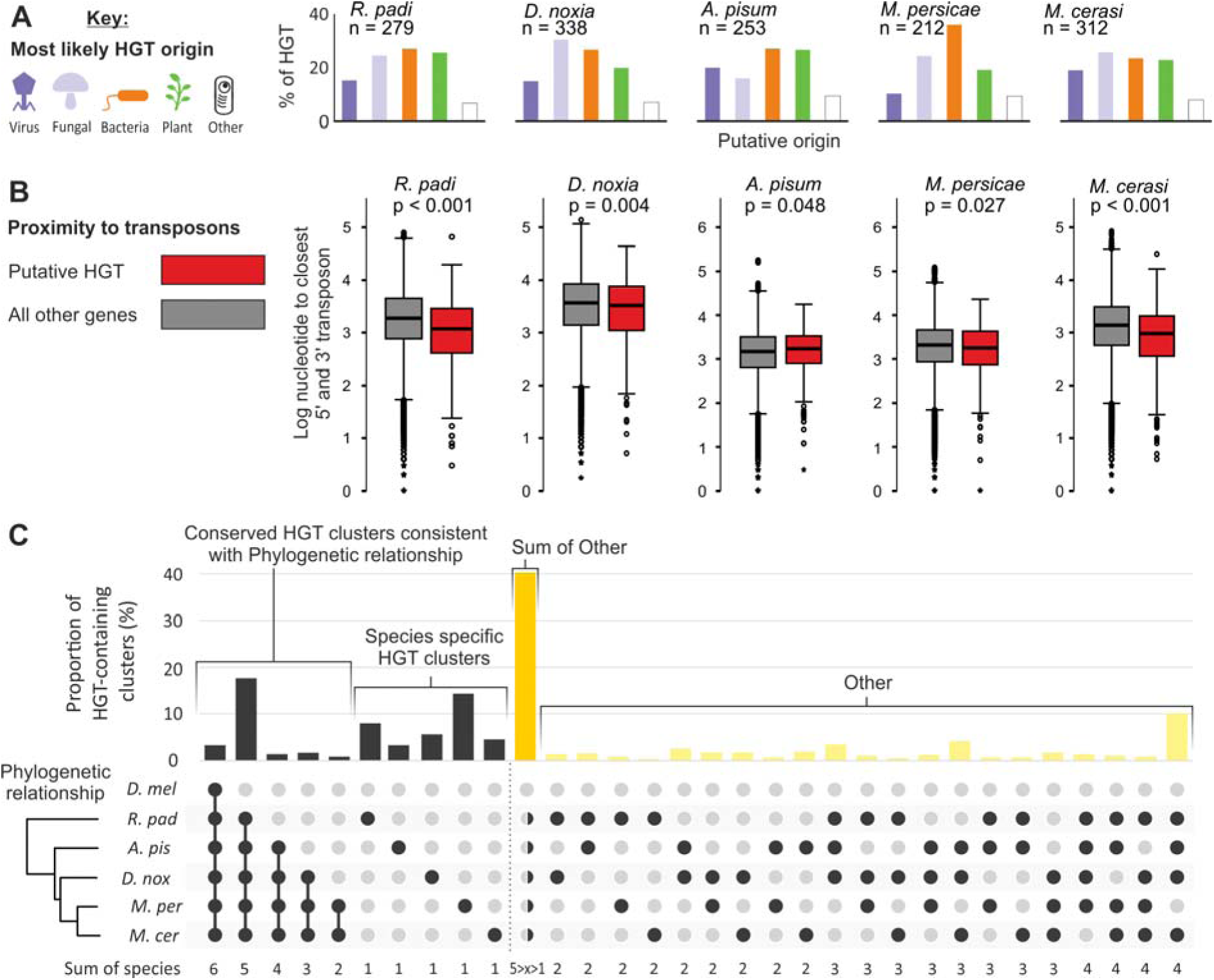
Putative horizontal gene transfer: origins and acquisition. We deployed a systematic genome-wide approach to identify putative horizontal gene transfer (HGT) events from non-metazoans, based on Alien Index calculations. **A**) The number of putative HGT events varies from 212 (*M. persicae*) to 338 (*D. noxia*). Histograms show the number of putative HGT events of viral (dark purple), fungal (light purple), bacterial (orange), plant (green), or non-metazoan Eukaryotes (white) for each aphid species. **B**) With the exception of *A. pisum*, putative HGT events (red) in aphid genomes are typically closer to their neighbouring 5’ and 3’ transposons than all other genes in the genome (grey). Mann-Whitney U test p-values range from 0.027 to <0.001. **C**) The histogram shows the proportion of HGT-containing clusters shared uniquely between the aphid species highlighted below. Approximately 40% of putative HGT-containing clusters are not consistent with the phylogenetic relationships of the different aphid species (dark yellow), but neither do they predominantly support any one other alternative (light yellow).

Intriguingly, predicted HGT events are generally closer to their 5’ and 3’ transposable elements when compared to the remainder of the genes in aphid genomes (Mann-Whitney U test, p-values range between 0.001 and 0.048, Figure 3B). This observation was similarly described for HGT events in a plant-parasitic nematode (reviewed in [28]), and perhaps is indicative of a general characteristic of HGT acquisition. Surprisingly, less than 20 % of predicted HGT loci are present in aphid-specific gene clusters (Figure 3C). This minority of HGT events likely originates before speciation and has been conserved to present day. Similarly, less than 15% of predicted HGT events are specific to a single aphid species, and likely originate after speciation events (Figure 3C). Remarkably, 40% of HGT containing clusters are not consistent with the phylogenetic positions of the different aphid species, but neither do they predominantly support any one other alternative (Figure 3C). Based on these observations we propose that most HGT events in aphids have complex evolutionary histories characterised by disparate gain and perhaps unsurprisingly frequent loss, and that HGT does not explain the large scale and recent gene birth.

Following transfer, predicted HGT events are apparently “normalized”. The distribution of AT content across HGT events is indistinguishable from that of the rest of the genes in the genome (Figure 4A). The AT content of HGT events from the species with the lowest average AT content (*M. cerasi*) is distinct from the AT content of the HGT events from the species with the highest average genomic AT content (*A. pisum*) (Figure 4B). Even HGT events predicted to be orthologous between *M. cerasi* and *A. pisum* have distinct AT content distributions that reflect their recipient “host” genome (Figure 4B). Finally, the variation in AT content of HGT events in *M. cerasi* only describes 10% of the variation of AT content in the corresponding *A. pisum* orthologous HGT events. This indicates that the DNA composition of the majority of HGT events in aphids match the composition of the recipient “host” genome rather than the “donor” non-metazoan genome. Despite the apparent normalization of DNA composition, the majority of HGT events have on average ~1 less intron per gene compared to the remainder of the genome (Mann-Whitney U test, p < 0.000 for all aphids tested). Nevertheless, the corresponding 5’ donor and 3’ acceptor splice sites are indistinguishable from the remainder of the genes in the aphid genomes, and are largely consistent with canonical CAG:GTAAGT (exon:intron) splicing (Figure 4C). The predicted *D. melanogaster* splice sites are in line with previous splice site predictions [29] [30].

**Figure 4.**
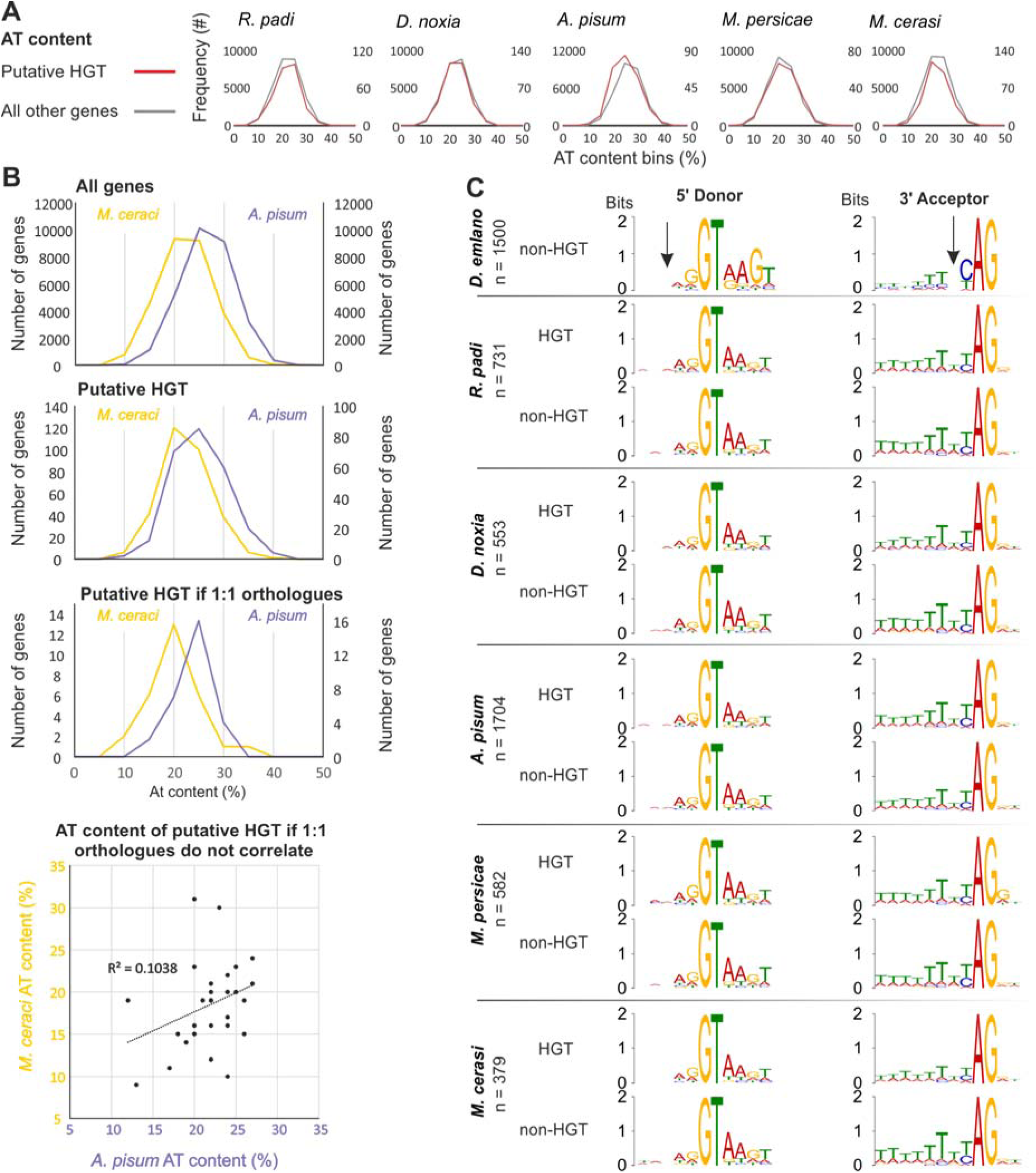
“Normalization” following horizontal gene transfer. **A**) The frequency distribution plots of AT content for putative HGT events (red) compared with T content of all other genes (grey) for *R. padi, D. noxia, A. pisum, M. persicae*, and *M. cerasi*. **B**) Comparison of AT content frequency distributions between *M. cerasi* (yellow) and *A. pisum* (purple) for all genes, putative HGT events, and putative HGT events that are putative 1:1 orthologues. **C**) Base composition of 5’ Donor and 3’ Acceptor splice-sites for a random selection of 1500 *D. melanogaster* genes is compared to that of all putative HGT events, and a randomly selected equal number of non-HGT evets, from *R. padi, D. noxia, A. pisum, M. persicae*, and *M. cerasi.* Black arrows indicate a consistent deviation from canonical splice sites.

The vast majority of putative HGT events have evidence of transcription, and the proportion of HGT events that have no measurable RNAseq expression is largely consistent with the remainder of the genome, with the notable exception of *A. pisum* (Additional file 6: Figure S3). The normalization of DNA composition of HGT events within the recipient genome in combination with evidence of transcription likely reflects the functional deployment following transfer and a contribution to the protein coding potential of aphid genomes. By comparing HGT predictions with a proteomics dataset that identified proteins present in saliva secretions [20] we are able to detect evidence of translation for a number of putative HGT events in *M. cerasi* (n=11) and *M. persicae* (n=3).

### The unusual genomic organisation and evolutionary history of predicted aphid effector repertoires

To compare aphid effector repertoires across the 5 different species, we (re)-predicted effector loci contained within the v1.0 annotations based on three modes of evidence, and as described previously [20]. In brief, we predicted effectors based on i) up-regulation in aphid head tissues (containing salivary glands) compared to aphid bodies without nymphs/heads in combination with the presence of signal peptide coding sequences (data set described by Thorpe et al., 2016), ii) presence in aphid saliva as determined by proteomics (data set described by Thorpe et al., 2016), and iii) similarity to previously described putative effectors [17] [16] [31]. Aphid genes with at least one mode of evidence were considered putative effector loci (Additional file 7: Table S4). These approaches were not applied to *D. noxia* due to the lack of tissue specific gene expression and saliva proteomics data.

In *R. padi*, *M. cerasi* and *M. persicae* approximately 2.4%, 2.7% and 3.6% of putative effectors are predicted to be acquired via HGT, approximately two and a half times the relative contribution to the remainder of the predicted proteome. One example in particular (Mca00616), is supported by saliva proteomics [20], and is part of a small cluster with *M. persicae* (Mpe23877), *D. noxia* (Dno04221) and *R. padi* (Rpa10410 and Rpa10411). The corresponding genes in *R. padi* are supported by saliva proteomics [20] but not predicted to be acquired via HGT (due to >70% protein identity to closest non-metazoan), the corresponding gene in *D. noxia* is predicted to be acquired via HGT, and the corresponding gene in *M. persicae* is predicted to be acquired via HGT, is supported by saliva proteomics [20], and is predicted to be an effector (Additional file 8: Figure S4).

Clustering only the putative effector gene content between aphid species (Figure 5A) revealed a different pattern to clustering the entire proteomes (Figure 2). Specifically, the pan-genome of putative aphid effector repertoires is dominated by singletons, with few highly connected clusters (Figure 5A). Given that most effectors were classified as multi-copy loci (Additional file 9: Figure S5), this suggests that while many effectors are part of paralogous/homolgous gene families within a species, often only one member of this family is predicted to be an effector. This can be a hallmark of either neofunctionalization following gene duplication, or loss of an effector gene following recognition by the plant immune system, and will need to be further explored.

**Figure 5.**
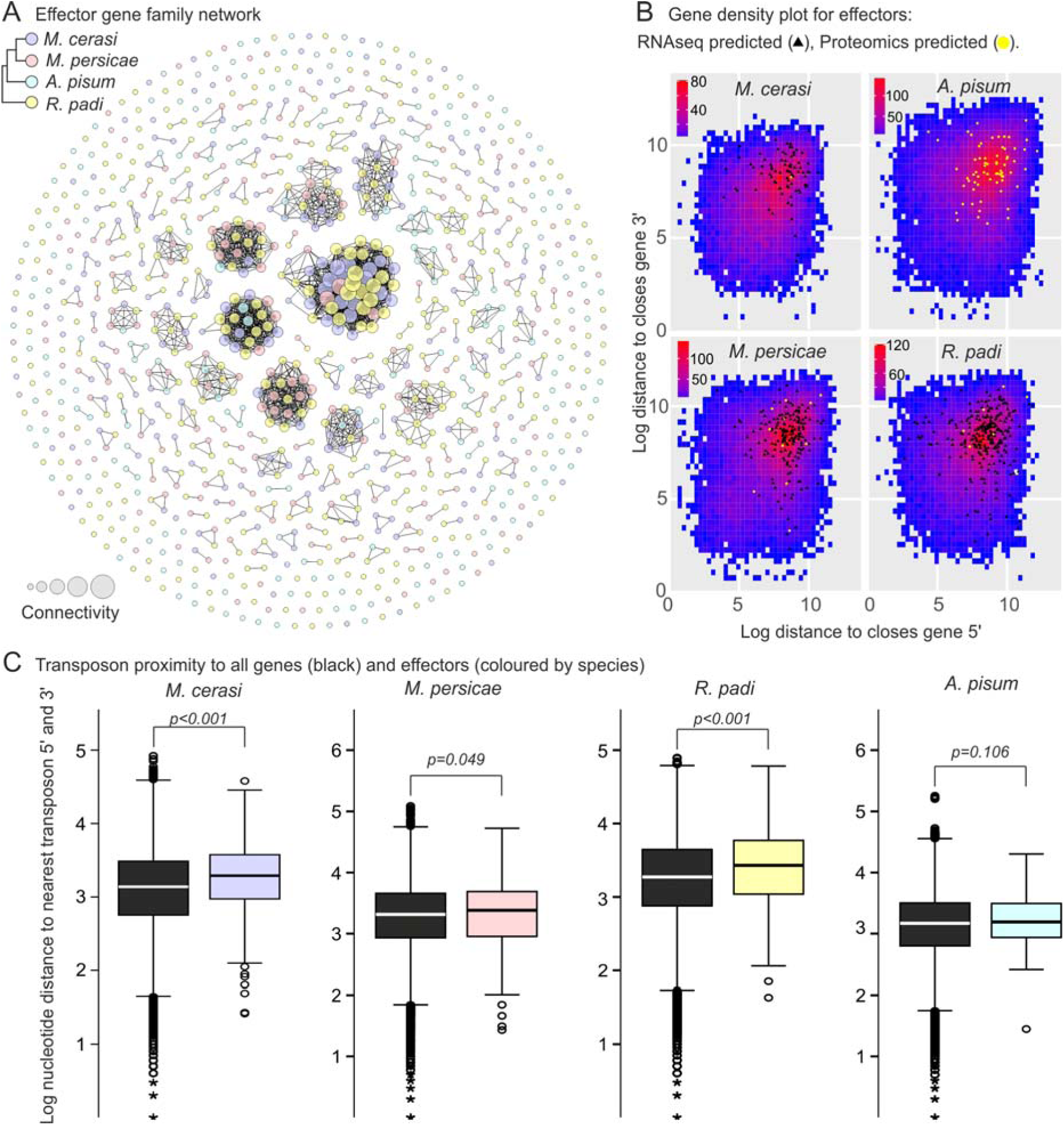
Effector repertoire and genomic organisation. **A**) Putative effector loci from all species were clustered with one another using BLAST. A network of sequence similarity was produced, where each node represents an individual effector locus, coloured by species. Connections between nodes indicate a minimum bitscore of 91, and node size is scaled by connectivity. **B**) The log nucleotide distance of each gene to its neighbour, 5’ (x axis) and 3’ (y axis) direction. Putative effectors are coloured by prediction method (RNAseq predicted – black triangle, Proteomics predicted – yellow circle). **C**) Box and whisker plots show the distance to nearest 5’ and 3’ transposable elements for putative effectors (coloured by species), and all other non-effectors (black). Distributions were compared using the Mann-Whitney U test.

Putative effectors are not randomly distributed across the aphid genomes, but are apparently partitioned into less gene-dense sub-domains (Figure 5B). Compared to all other genes in the aphid genomes, putative effectors are significantly further from their neighbouring genes in both the 3’ and 5’ prime directions (Mann Whitney U-test, p<0.000, Figure 5B). Similarly, effectors from distinct eukaryotic plant pathogens are often located in less-gene dense regions within the genomes [32] [21] [33]. However, the classical signature of a close genetic association of putative effectors and transposable elements, as reported for oomycetes, nematodes and fungi, does not manifest in the aphid genomes [32] [21] [33]. With the exception of *A. pisum*, putative effectors are actually further from their nearest transposable element in both the 3’ and the 5’ prime direction when compared to the remainder of the genes in the genome (p<0.01 and p<0.05, respectively. Mann Whitney U-test, Figure 5C).

The presence of effectors in less-gene dense regions of the genome is hypothesised to coincide with regions of high mutability, thus providing a means for rapid evolution of genes under high selection pressure from the plant host immune system [22]. Consistent with this, we identified 30 orthologous gene clusters, containing 170 putative effectors, as being under diversifying selection (DN/DS greater than 1.0 (Additional file 10: Table S5), notably including well-characterized effectors C002 (DN/DS = 2.35) and Me10-like (DN/DS = 1.60) [20].

### Genetic linkage, and shared transcriptional control, of a subset of predicted aphid effectors

We initially noted that putative orthologues of the previously characterised effectors Me10 [19] and Mp1 [17] [34] [14] (here referred to as *Me10-like* and *Rp1*) are tightly genetically linked in the *R. padi* genome. These two genes are present in a head to tail orientation, 5417 bp from the end of the first (*Rp1*, *Rpa14995*) to the start of the last (*Me10-like*, *Rpa14996*). This same genetic linkage, and an identical genomic organisation, is conserved in all 5 aphid species (Figure 6A). Remarkably, genes and transposons adjacent to this effector pair are different in every aphid species, indicating a lack of synteny in the corresponding genomic regions. Effector gene co-location does appear to be a feature of effectors, albeit not universal: effector *COO2* is present in a large syntenic block of non-effector loci conserved in all aphids (Figure 6B), while 25.8% of *R. padi* putative effectors have another putative effector as an adjacent genomic neighbour (c.f. ~3% expected by chance). We observed that the promoter region of the 5’ gene of each of the *Mp1-Me10-like* pair is highly similar, and may be indicative of shared transcriptional control (Additional file 11: Figure S6).

**Figure 6.**
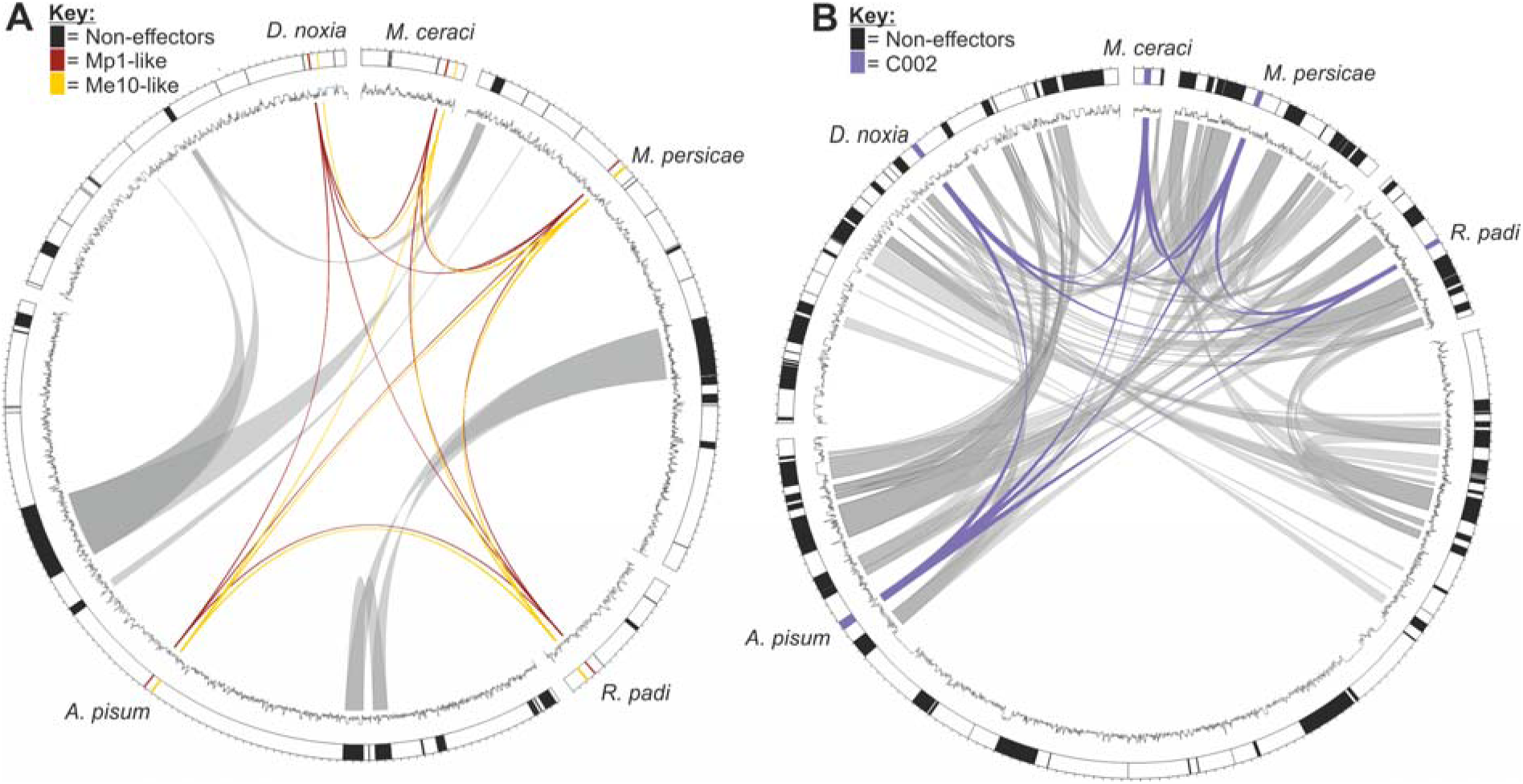
Tight genomic association of *Mp1* and *Me10-like* effectors. Circos plots showing selected scaffolds/contigs of the *R. padi, D. noxia, A. pisum, M. persicae*, and *M. cerasi* genome assemblies **A**) The characterised effectors Mp1-like and Me10-like are conserved as an adjacent pair in all species. Every neighbouring gene and transposon is different, in all genome assemblies. **B**) The characterised effector C002-like is present in a syntenic block that is largely conserved in all species.

To determine whether *Me10-like* and *Rp1* are under shared transcriptional control, and to what extent transcriptional plasticity contributes to aphid interactions with host versus non-host plant species, we sequenced the transcriptome of *R. padi* and *M. persicae* after feeding on an artificial diet for 3 or 24 hours, a host plant for 3 or 24 hours, and a non-host plant for 3 or 24 hours (each with five replicates prepared in environment controlled growth cabinets, conducted at the same time of day). *R. padi* shows an extremely inconsistent transcriptional responses to these quite different stimuli, such that the response is more highly correlated between samples than within replicates (Additional file 12: Figure S7 and Additional file 13: Table S6). Nevertheless, expression of the *R. padi Rp1*/*Me10-like* gene pair was almost perfectly correlated (Figure 7A): measuring variation in the expression of *Me10-like* describes 99% of the variation in expression of *Rp1* (R^2^ 0.99, Figure 7A). No such correlation was observed when comparing the expression of *Rp1* with its adjacent non-effector gene in the opposite direction (Rpa14994, R^2^ 0.06, Figure 7A). We identified 5 other pairs of effector genes that are adjacent in all aphid species, but their expression did not correlate to the same extend as the *Rp1*/*Me10-like* pair (Additional file 14: Figure S8), and these did not necessarily share the same orientation. The fact that the *Rp1*/*Me10-like* genetic linkage has persisted throughout evolution in spite of considerable local re-arrangements, coupled with shared transcriptional control, is strongly indicative of functional linkage.

**Figure 7.**
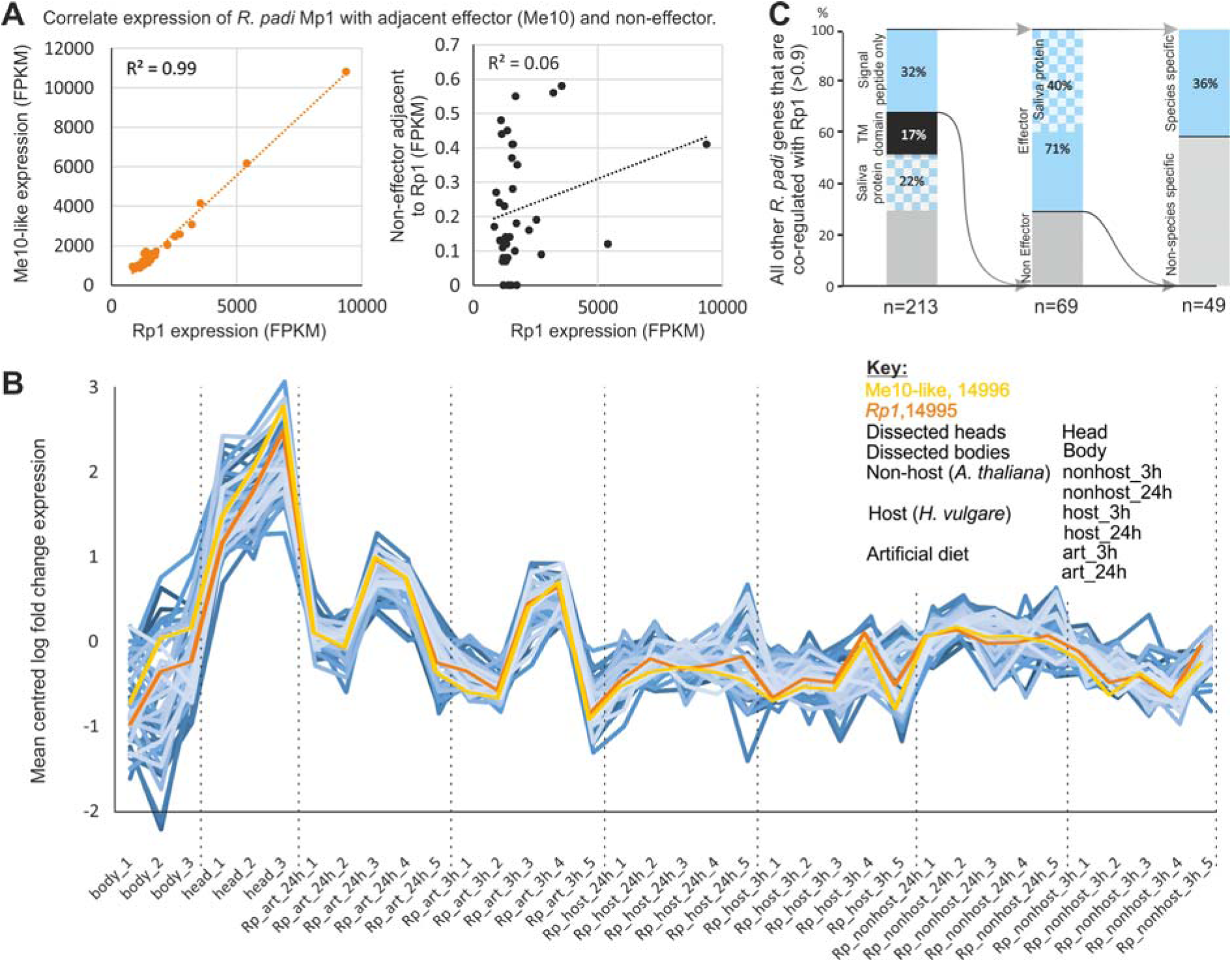
Shared transcriptional control of a subset of the effector repertoire. **A**) Correlating the normalized RNAseq expression of the *Rp1* effector with its adjacent effector *Me10-like* (orange) reveals almost perfect concerted expression across a range of diverse stimuli (Figure S7). No such correlation is observed with the adjacent non-effector (black). **B and C**) Identification of all other genes in the *R. padi* genome that are co-regulated with the Rp1:Me10-like pair based on a >90% Pearson’s correlation (blue, n = 213). Of these, 32% are secreted (n = 69). Of these, 71% are predicted to be effectors (n = 49). Of these, 36% are present in MCL clusters that exclude all other aphid species (n = 18). Remarkably, of the 144 genes co-regulated with the *Rp1:Me10-like* pair based on a >90% Pearson’s correlation that do not harbour a signal peptide sequence indicative of secretion, 46 were previously detected in the saliva of *R. padi* by mass spectrometry (blue and white checker box) [20].

Using the genetically linked *R. padi Rp1/Me10-like* effector pair, we sought to identify other genes that are similarly transcriptionally, but not physically, linked, in the *R. padi* genome. Despite the apparently inconsistent transcriptional response to stimuli, there are 213 other loci in the *R. padi* genome that mirror this response with a Pearson’s correlation of > 90% (Figure 7B). Of these 213, 32% were predicted to encode secretory proteins (a 4.5 fold enrichment over the remainder of the genome). Of these 69, 71% were already predicted to be effectors (n=49, Figure 7C). Of these 49 effectors, 36% were present in gene clusters specific to *R. padi* (n=18). Taken together, this suggests that with just two criteria, 1) concerted expression with a highly conserved effector pair and, 2) the presence of a signal peptide for secretion, a 71% accuracy of effector identification can be achieved. These predictions work similarly, albeit to a lesser extent in *M. cerasi* (32% were predicted secreted, 62.5% of which were already predicted to be effectors), and *M. persicae* (57% were predicted secreted, 16% of which are already predicted to be effectors). We did not have sufficient data to produce similar outputs for *A. pisum* and *D. noxia*. Remarkably, of the 213 that correlate >90% with the *Rp1:Me10-like* pair that are not predicted to encode a secretion signal, 46 have been detected in the saliva of *R. padi* using a proteomics approach [20]. This is a substantial proportion of all proteins detected in the saliva of *R. padi* (30%), is numerically more than those with a classical signal peptide for secretion, and may question the suitability of canonical secretory protein prediction pipelines. Concerted expression of effectors has been reported in a plant pathogenic fungus, and likely relies on an epigenetic control mechanism [35]. Whether epigenetic control also is responsible for the tight co-regulation of a significant subset of aphid effectors remains to be elucidated.

## Conclusions

Aphid species are highly diverse with regards to many phenotypic and ecological traits, which are defined by unknown mechanisms of genome evolution. In this study we reveal a complex history of ancient gene duplication and relatively recent gene birth in aphids, as well as contribution of HGT events, and disparate gain and loss of genes, to the protein-coding content, and effector complement, of aphid species. We identified several aphid effector pairs that are genetically linked across aphid species, one of which also showed tight co-regulation of transcription with a substantial subset of putative effectors. Exploiting transcriptional linkage for utility, we develop a series of criteria to expand the putative effector repertoire of aphids, and potentially implicate non-classical secretion in aphid parasitism.

## Methods

All data are available under accession numbers PRJEB24287, PRJEB24204, PRJEB24338 and PRJEB24317. Assembled genomes and gene calls are available at http://bipaa.genouest.org/is/aphidbase/ and http://mealybug.org. All custom python scripts used to analyse the data use Biopython [36], are available on Github, and are cited in the text where appropriate.

### Aphids stocks and material

Aphids were maintained in growth rooms at 18 °C with a 16 h light and 8 h dark period. *M. persicae* (genotype O) was maintained on oil seed rape, a clonal line of *M. cerasi* was maintained on American Land Cress (*Barbarea verna*) and a clonal line of *R. padi* was maintained on barley (*Hordeum vulgare* cv. *Optic*).

### DNA extraction and sequencing

Aphids were collected and subjected to one ethanol wash, with agitation, to help remove fungal and bacterial contamination, followed by three sterile distilled water washes. DNA was extracted using Qiagen Blood Tissue extraction kit following manufacturer’s protocol, followed by a DNA ethanol precipitation step to improve DNA purity. DNA quality was assessed using a Nanodrop (Thermo Scientific) prior to sending the Earlham Institute, Norwich, for PCR-free library preparation and sequencing (insert size ~395bp). Illumina-HiSeq 2X250bp (and 2X150bp for *M. c*erasi) paired-end sequencing was performed.

### Filtering, quality control and genome assembly

The raw reads were assessed for quality before and after trimming using FastQC [37]. For quality control the raw reads were quality trimmed using Trimmomatic (minimum phred Q15) [38]. An iterative process of assembly and contaminant removal was performed. For early iterations of the assembly, CLC (version 4.1.0) was used due to rapid assembly and coverage mapping. To remove contaminant reads, the assembly was compared to the non-redundant database (nt) using BLASTn (megablast), the assembly was also searched against the genome sequence of *A. pisum* to facilitate the identification of Arthropoda contigs, SWISS-Prot database and GenBank NR using DIAMOND (v0.7.9.58) in sensitive mode [39]. The DIAMOND-BLAST vs NR data was taxonomically annotated using https://github.com/peterthorpe5/public_scripts/tree/master/Diamond_BLAST_add_taxonomic_info. The resulting taxonomically annotated BLAST results and genomic read coverage generated by CLC (mapper) were used as input to Blobtools [40]. Reads that contributed to the assembly of contigs similar to bacteria, fungal or virus sequences were removed in an iterative approach using Mirabait *K*=99 [41]. This was repeated 8 times for *M. cerasi* and 5 for *R. padi.*

The final “cleaned” datasets were converted from .fastq to .bam files using custom python scripts and were assembled using DISCOVAR [42]. All assemblies were assessed for “completeness” using CEGMA [24] and BUSCO using Arthropoda Hidden Markov Models (HMM) [23]. Statistics on genome assemblies were generated using: https://github.com/sujaikumar/assemblage/blob/master/scaffold_stats.pl. All scripts and commands used for genome assembly are available under the https://github.com/peterthorpe5/Methods_M.cerasi_R.padi_genome_assembly

### Gene prediction and annotation

Due to a lack of publically available known genes from both *M. cerasi* and *R. padi*, the approach of using known sequences to train MAKER [43] was not used. A preliminary approach was taken. Augustus [44] gene prediction, using RNAseq hints for each species was performed using the “Pea_aphid species config files” bundled with Augustus [45] (Gene models: v0.9). RNAseq was mapped to the genomes using splice aware aligner STAR [46] allowing a maximum of 7 mismatches, RNAseq intron hints was generated with bam2hints (a script bundled with Augustus). Additional RNAseq data for each species was obtained from: *R. padi, M. persicae, M. cerasi* (PRJEB24317) and PRJEB9912 [20]*, A. pisum* (unpublished) and PRJNA209321 [45], *D. noxia* SRR1999270 [4]. Once a set of gene models were predicted, the RNAseq for each species was mapped back to the nucleotide CDS gene prediction (exome) of that species using SNAP [47], to determine the percentage of RNAseq that maps. This did not allow reads that spanned the start and stop codon, and SNAP is a DNA aligner, thus spliced reads resulted in a lower Q mapping score. All mapping was performed in the same way for comparison purposes only. Predicted proteins were DIAMOND-BLASTp [39] searched against NR, and taxonomically annotated as described above. Due to poor RNAseq mapping results, the “Pea_aphid config files” guided gene models were deemed unsatisfactory (see results). Therefore an alternative, *de novo*, approach was taken.

The final gene models for all species were predicted using BRAKER (version 1.8) [48] and intron RNAseq guided hints (see above) (Gene models: v1.0). BRAKER uses Genemark-ET [49], with the RNAseq hints and Eukaryote HMM models to predict genes and retrain Augustus. Trained Augustus was used in conjunction with RNAseq intron hints to predict gene models v1.0. Gene models were annotated using Blast2GO version 2.8, database September 2015 [50], Interproscan [51], PFAM [52], DIAMOND-BLASTp versus NR [39]. The DIAMOND BLAST output was taxonomically annotated as mentioned above. BLAST output was taxonomically filtered to remove Pea aphid “hit” using: https://github.com/peterthorpe5/public_scripts/blob/master/blast_output/top_BLAST_hit_filter_out_tax_id.py

### Endosymbiont genome assembly

To assemble the Buchnera spp. genome from the genomic data, raw reads were trimmed of adapter sequences and low quality bases (Phred <30), and assembled using SPAdes (version 3.5) using k=77,99,127 [53]. From this assembly, one of the contigs corresponded to the expected genome size of the endosymbiont, and shared considerable sequence similarity to other Buchnera genomes. The Buchnera spp. genomes were annotated with the web-server instance of RAST [54].

### Transposon-like sequence prediction and repeat masking

To predict transposons and repetitive regions, an aphid specific database was generated using RepeatModeller (version 1.0.8) [55]. The database was classified using Censor [56]. Repeatmasker (version 4.0.6) [57] using this classified database and Repbase was used to identify repetitive regions and transposons. LTRharvest (genometools-1.5.8) [58] and TransposonPSI (version 08222010) [59] were also used to identify transposons. A consensus prediction was generated and .gff formatted. https://github.com/HullUni-bioinformatics/TE-search-tools. Transposon and gene distances were calculated using: https://github.com/peterthorpe5/public_scripts/tree/master/transposon_analysis

### Alien index - Detection of horizontal gene transfer events and putative contamination

To detect candidate horizontal gene transfers (HGT) events, an Alien Index (AI) was calculated as described in [25, 26]. All predicted proteins were compared to NR using DIAMOND-BLASTp, with kingdom and tax_id assignment, and an e-value threshold of 1e^−5^. An AI could only be calculated for a protein returning at least one hit in either a metazoan and non-metazoan species, as stated in the following formula: AI = log((Best Evalue for Metazoa) + e-200) - log((Best E-value for Non-Metazoa) + e-200)

When neither metazoan nor non-metazoan BLAST results were identified, the query sequence was removed from downstream analysis. BLAST results in the phylum Arthropoda (which the aphids of interest belong) were ignored for the calculation of AI, to allow detection of HGT events that may be shared with other related species. An AI > 30 corresponds to a difference of magnitude e^10^ between the best non-metazoan and best metazoan e-values and is estimated to be indicative of a potential HGT event [25].

Sequences with an AI> 30 and >70% identity to a non-metazoan sequence were considered putative contaminants and removed from further analyses (Additional file 15: Table S7). Horizontal gene transfer prediction tool set is available at Github: https://github.com/peterthorpe5/public_scripts/tree/master/Lateral_gene_transfer_prediction_tool. Intron splice sites were extracted using: https://github.com/DRL/GenomeBiology2016_globodera_rostochiensis/tree/master/scripts and log plots were generated using MEMEsuite [60]. Metabolic pathways were predicted using the entire predicted proteome of each species using the KEGG Automatic Annotation Server [61].

### Transcriptomic analyses upon aphid exposure to host, non-host plants and artificial diets

To determine the extent transcriptional plasticity contributes to aphid interactions with host and nonhost plants, we sequenced the transcriptome of *R. padi* and *M. persicae* after feeding on an artificial diet for 3 or 24 hours, a host plant for 3 or 24 hours, and a non-host plant for 3 or 24 hours. For *R. padi*, barley is considered a host and Arabidopsis is considered non-host [62]. For *M. persicae*, Arabidopsis is considered host and barley is considered non-host [63]. For both species, the artificial diet consisted of 15 % sucrose, 100 mM L-serine, 100 mM L-methionine and 100 mM L-aspartic acid with a pH of 7.2 (KOH) [64].

Barley plants (cv Optic) were pre-germinated in Petri dishes with wet filter paper for three days in the dark. Plants were moved to a growth room and grown for 7 days prior to aphid infestation. Arabidopsis plants were sown directly in soil and grown for 5 weeks prior to aphid infestation. Artificial diets were prepared and placed between Parafilm sheets according to [20]. Plant growth as well as aphid exposure to plant and diet were carried out under 8 hours of light (125 μmol photons/m^2^.s), at 22 °C and 70% humidity.

For transfer of *R. padi* and *M. persicae* aphids from stock plants to barley and Arabidopsis, 15 mixed-aged apterous aphids were enclosed in a single clip cage, with one clip cage per plant, and 6 plants per plant-aphid combination per time point (3h and 24h). The clip cage was placed in the middle of the 1^st^ leaf for barley, and covering 1-2 fully expanded leaves for Arabidopsis. For the artificial diet treatment, 100 mixed-aged apterous aphids were used per time point in a single artificial diet container. All aphids were collected 3h and 24h after exposure to plants or diet and flash frozen in liquid nitrogen, and aphids from the 6 individual plants per plant-aphid combination per time point were pooled into one single tube. In total, 5 independent biological replicates were performed of the whole experiment. Individual replicates were set up at the same time of day to avoid variability due to the aphid or plant circadian cycle. Replicates of host and non-host plants treatments were set in different weeks over a 2 month period at approximately 9:00h, with the 3h time point collected at 12:00h the same day, and the 24h time point at approximately 9:00h the next day. Artificial diet treatments were not set-up in parallel to the plant treatments, but on consecutive days, between 10:00 and 12:00h, with collection of the 3h time point occurring between 13:00 to 13:30h the same day, and collection of the 24h time point between 11:00 to 12:00h the next day.

RNA was extracted from 70-90 aphids with the Qiagen RNeasy Plant Mini Kit^®^ following the manufacturer’s protocol. RNA quality was assessed using agarose gel electrophoresis and the Agilent 2100 Bioanalyzer. Approximately, 2.5 μg of total RNA per sample (60 samples total) was submitted to TGAC (The Genome Analysis Centre, Norwich Research Park) for Illumina TrueSeq library preparation and sequencing (100 bp paired end).

Temporal RNAseq data described above were analysed with spatial RNAseq data of a previous study (PRJEB9912 [20]). All raw RNAseq reads were assessed using FastQC [37], and low quality bases were removed using Trimmomatic (Minimum Phred score 22) [38]. Reads were mapped to the corresponding genome using STAR version2.5.1b [46].

The resulting bam file was assembled using Trinity (version 2.1.1) [65]. The assembly was subjected to quality control using Transrate [66]. Transcript abundance was quantified using Kallisto [67]. Differential expression analysis was conducted using EdgeR [68], using minimum threshold of Log2 fold change and a FDR p<0.001. Coding sequence from transcripts was predicted using TransDecoder [65].

### Effector identification and comparisons

To compare aphid effector repertoires across the 5 different species, we (re)-predicted effector loci contained within the v1.0 annotations based on three modes of evidence, and as described previously [20]. In brief, we predicted effectors based on i) up-regulation in aphid head tissues (containing salivary glands) compared to aphid bodies without heads in combination with the presence of signal peptide coding sequences (data set described by Thorpe et al., 2016), ii) presence in aphid saliva as determined by proteomics (data set described by [20]), and iii) similarity to previously described putative effectors [17] [16] [31]. Aphid genes with at least one mode of evidence were considered putative effector loci (Additional file 7: Table S4). These approaches were not applied to *D. noxia* due to the lack of tissue specific gene expression and saliva proteomics data. The effector repertoire network of all species was generated by calculating the BLASTp bit score of pairwise comparisons between effectors of all species. An array of pairwise bit scores was parsed to gefx format using a custom python script (https://github.com/sebastianevda/SEvdA_Gephi_array_to_gefx), and visualised using Gephi [69].

### Promoter analyses

The genomic 5’ region to genes of interest was obtained using custom python script https://github.com/peterthorpe5/public_scripts/tree/master/genomic_upstream_regions. Motif enrichment was performed using the differential motif discovery algorithm HOMER [70].

### Comparative genomics

An MCL all vs all network was generated using the predicted proteomes of *R. padi, D. noxia, A. pisum, M. persicae, M. cerasi*, and the outgroup model insect *D. melanogaster.* Similarity was assessed using DIAMOND BLASTp (1e-31) and clustered using MCL (inflation value of 6). All MCL analyses were performed using Biolinux 7 [71]. Individual sequence alignments were carried out using Muscle v3.8.31 [72], and visualised using the BoxShade web server (https://www.ch.embnet.org/software/BOX_form.html)

### Gene duplication and synteny analyses

Gene duplication and synteny analysis was performed using the similarity searches from DIAMOND-BLASTp (evalue 1e-5) with MCSanX toolkit [73]. Synteny between scaffolds was visualised using Circos 0.67-7 [74].

### Phylogenetic inference

Single copy orthologous genes were identified using in all five aphid species studies, and the out-group *D. melanogaster*. Only those sequences identified in all genome assemblies, and classified as single copy loci in all assemblies, were studied (n=386). For a given BUSCO gene in a given species, if the gene length deviated by more than 5% from the average for that BUSCO gene in all other species, that BUSCO gene was not analysed further for any species (remaining n=123). The amino acid sequences of the remaining 123 highly conserved BUSCO genes were aligned and refined using MUSCLE (Additional file 16). Individual BUSCO alignment were concatenated and a partition file generated using a custom python script (https://github.com/sebastianevda/SEvdA_Gephi_array_to_gefx/blob/master/cat_alignments_rename_names_write_partition_file.py). Model selection for each partition, and phylogenetic inference, was carried out using the IQ-TREE webserver to generate a consensus tree of 1000 bootstraps [75].

### DN/DS analysis

A 1:1 Reciprocal Best BLAST Hit network was generated from the predicted amino acid sequences, using a minimum threshold of 70% identity and 50% query coverage [76] and clustered using MCL (version 12-135) [77] with an inflation value of 6. The number of species contained in a cluster was obtained using mcl_to_cafe.py [78]. DN/DS values for each cluster that contained a predicted effector was calculated. Within each cluster, deduced protein sequences were aligned using MUSCLE (version 3.8.31) [72], and the nucleotide sequences back-translated onto the alignments (https://github.com/peterjc/pico_galaxy/tree/master/tools/align_back_trans) [36]. Alignments were manually curated using Jalview [79] by removing non-consensus, possibly miss-predicted 5’ and 3’ regions. Modified alignments were subjected to DN/NS analysis using CodonPhyml (version 1.0) [80].

## Additional Data Files

**Additional file 1: Figure S1.**
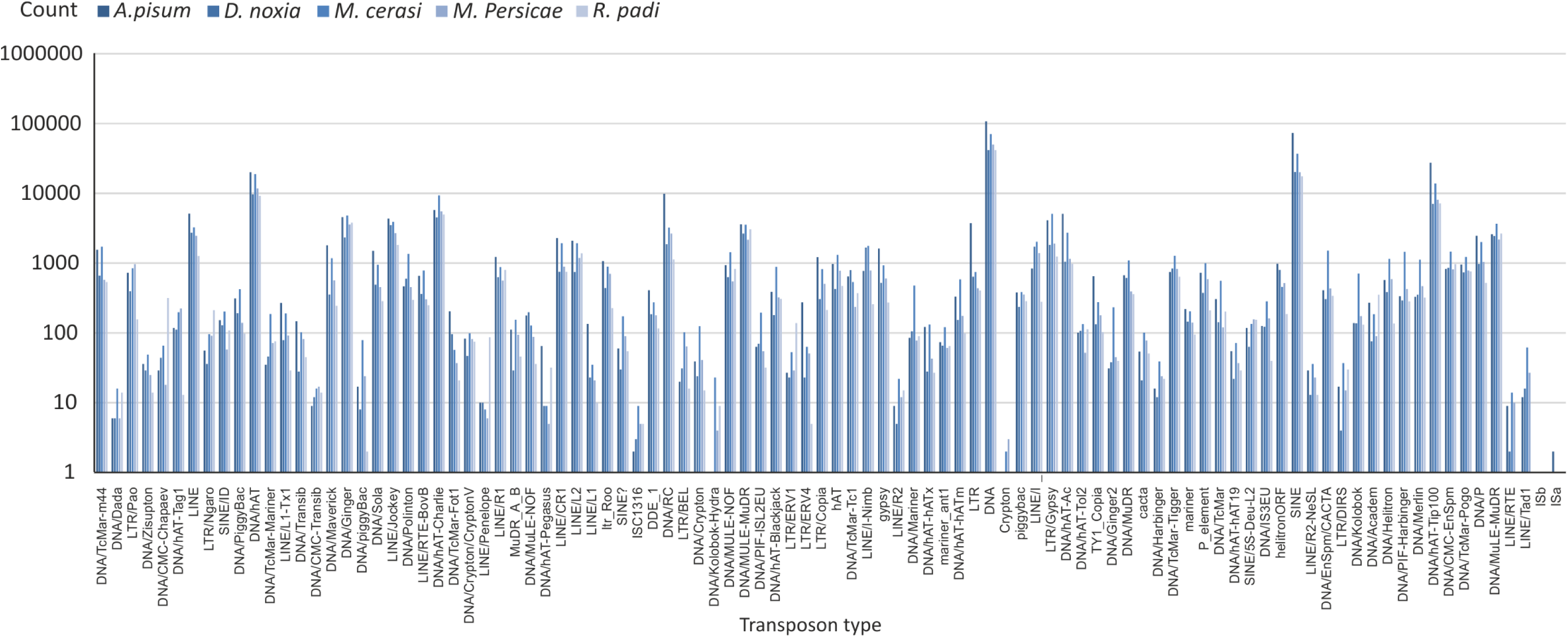
Transposable element prediction in aphid genomes. The number and types of transposable element-like sequences present in the genomes of *A. pisum, D. noxia, M. cerasi, M. persicae*, and *R. padi*. Counts are displayed on a per transposon type basis for each species. All aphids tested have very similar absolute transposon numbers, and very similar relative transposon composition, with few exceptions (e.g. Crypton, ISa, and ISb).

Additional file 2: Table S1 - Gene model statistics v0.9 and v1.0

Additional file 3: Table S2 - Gene duplication categories

Additional file 4: Table S3 - Putative Horizontal Gene Transfer events

**Additional file 5: Figure S2.**
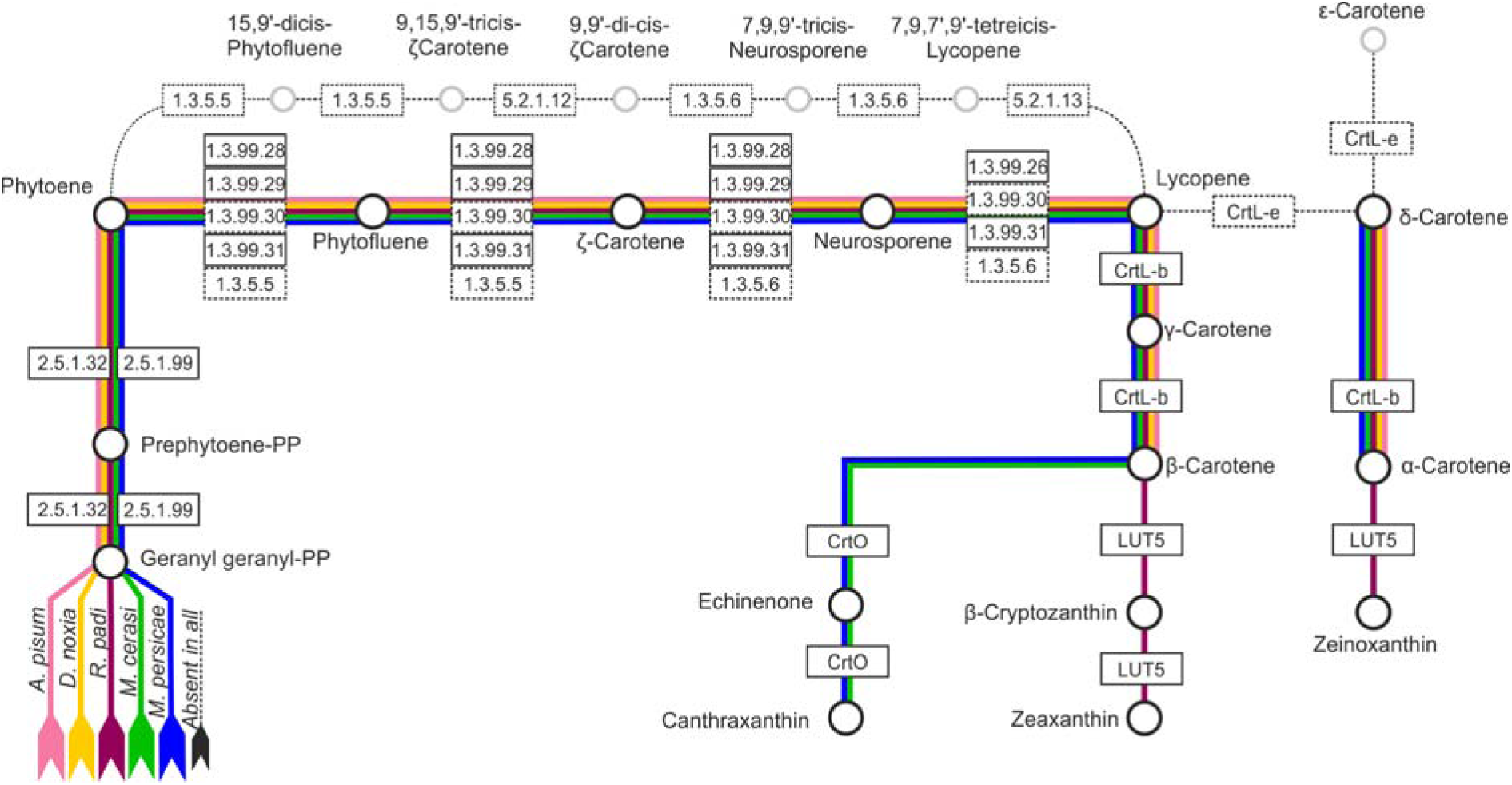
Metabolic pathway analysis for carotenoid biosynthesis. Metabolic pathway analysis using the KEGG Automatic Annotation Server [61]. Enzymes are represented by boxes (EC codes/name within), while products/substrates are represented by circles. Lines which link circles indicate the pathway/s that each species is theoretically capable of carrying out based on the predicted proteome. Lines are coloured by species: *A. pisum*, pink; *D. noxia*, yellow; *R. padi*, purple; *M. cerasi*, green; *M. persicae*, blue. Pathways that are apparently absent in all species are represented by black broken lines, where boxes and circles are greyed. Uniquely, the *R. padi* genome apparently lacks the Phytoene desaturases, and encodes LUT5. Only the Myzus genomes appear to encode CrtO. The *R. padi* gene predictions apparently lack the phytoene desaturases, but these EC classes are encoded by the *de novo* transcriptome for this species.

**Additional file 6: Figure S3.**
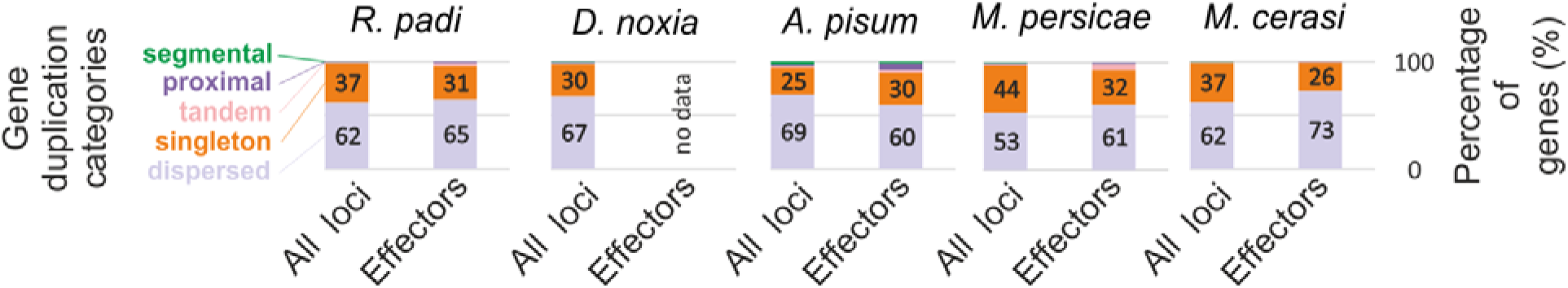
Putative HGT events with evidence of transcription. Comparison between the expression of putative HGT (black) events and all other loci (white). The proportion of genes with no evidence of transcription is plotted on the y axis.

Additional file 7: Table S4 - Putative effector loci

**Additional file 8: Figure S4.**
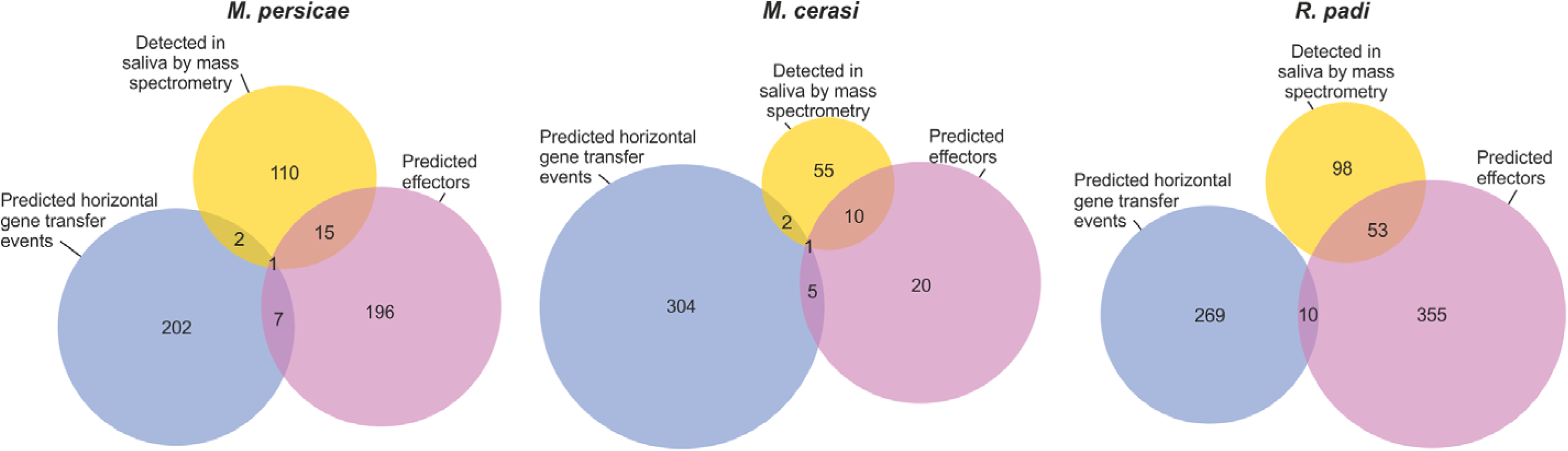
Horizontal gene transfer has contributed to the putative effector repertoires of aphids. For *M. persicae, M. cerasi* and *R. padi*, the overlap between putative effectors loci (pink), loci putatively acquired by horizontal gene transfer (blue), and loci encoding proteins detected in the salivary secretions as determined in [20].

**Additional file 9: Figure S5.**
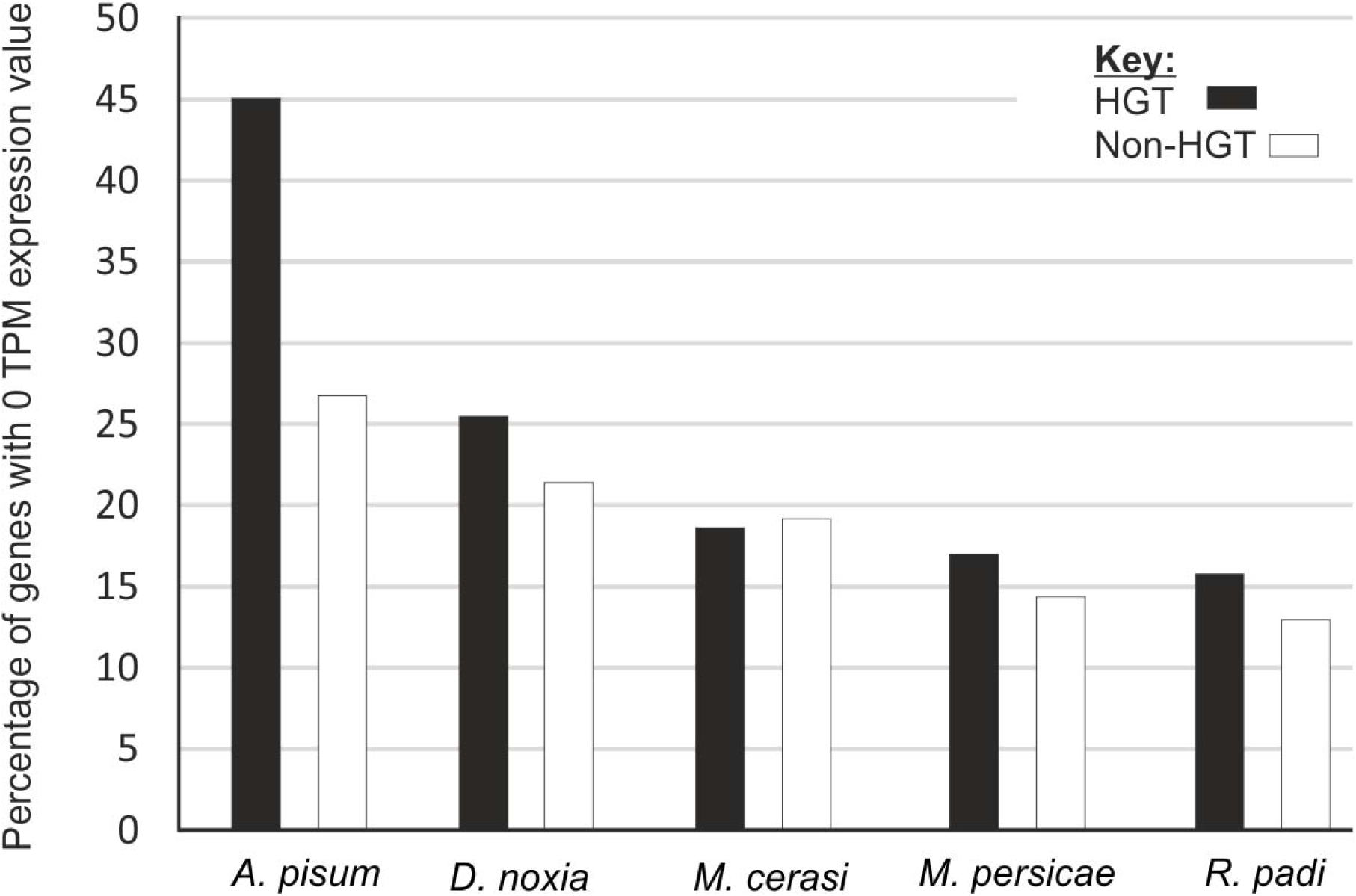
Gene duplication categories of all loci compared to putative effectors. Gene duplication categories on a per species basis, for *R. padi, D. noxia, A. pisum, M. persicae*, and *M. cerasi*. Internal numbers show the percentage of dispersed (light purple) and singleton (orange) categories. For each species, the proportion of gene duplication categories across all genes, or only the effectors, are shown.

Additional file 10: Table S5 - Putative effectors under positive selection

**Additional file 11: Figure S6.**
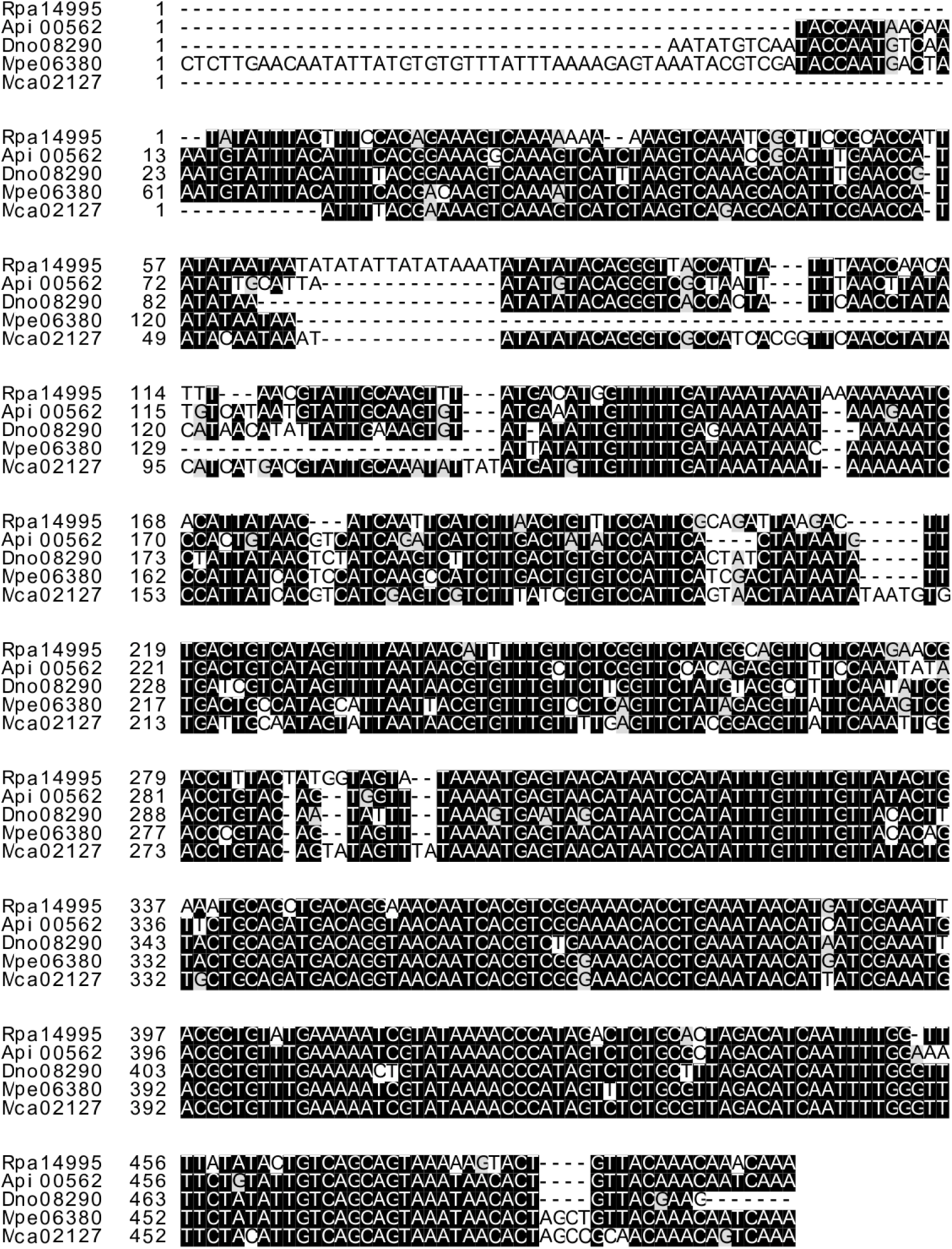
Promoter alignment of *Mp1-like* from each *Mp1-Me10* pair. Multiple sequence alignment of approximately 500 bp 5’ of the first gene (*Mp1-like*) in the adjacent *Mp1-like/Me10-like* pair from *R. padi, D. noxia, A. pisum, M. persicae*, and *M. cerasi*. Black boxes indicate identical nucleotides in for those species highlighted.

**Additional file 12: Figure S7.**
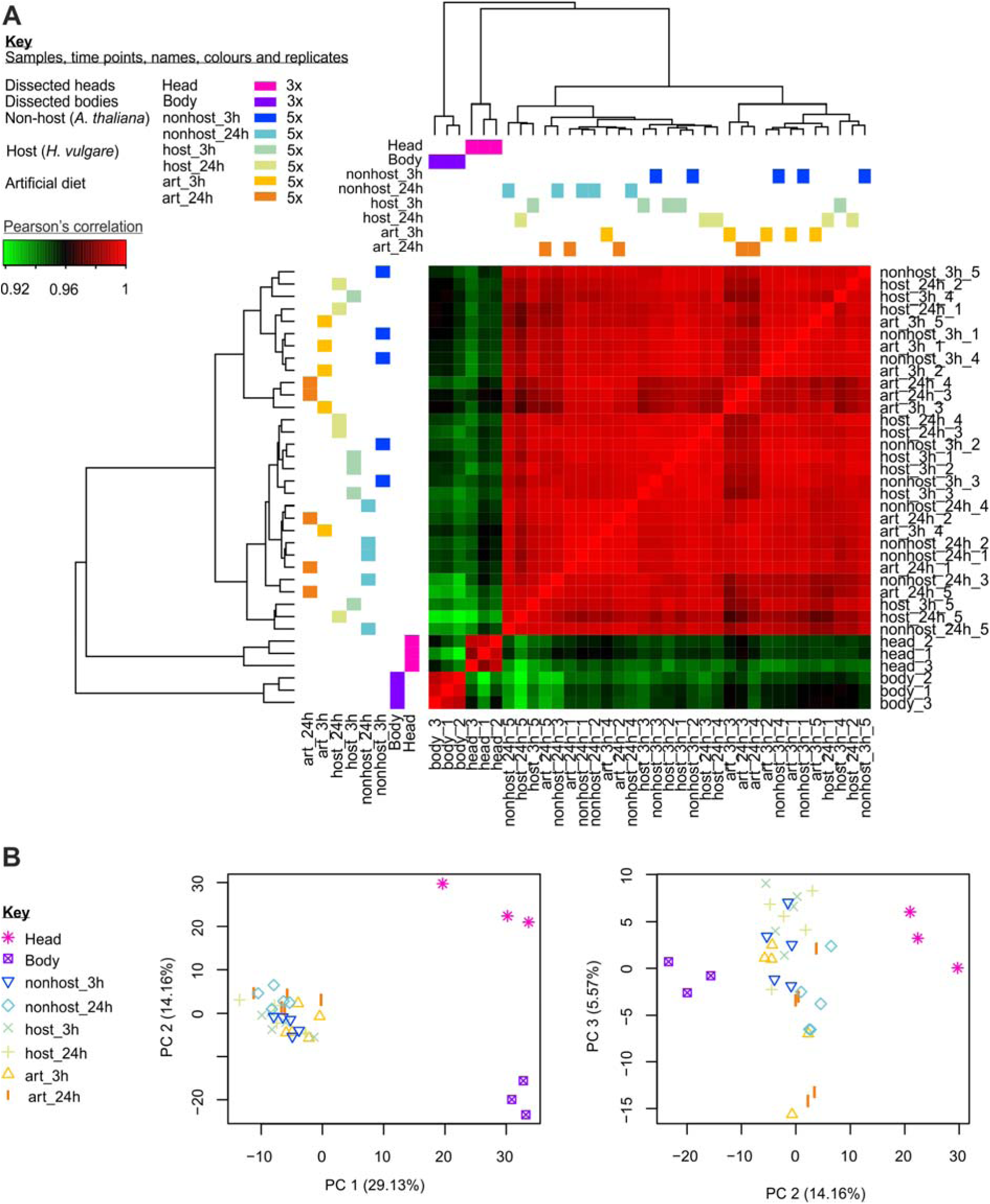
Inconsistent transcriptional response to artificial diet, host, and non-host. Genome-wide analysis of *R. padi* transcriptional responses to a range of stimuli (artificial diet (art), feeding on host (*H. vulgare*), and feeding on a non-host (*A. thaliana*) for 3 and 24 hours), and comparison to previously published tissue-specific transcriptome of dissected heads and bodies [20]. **A**) Clustering of transcriptional response reveals that *R. padi* exhibits an extremely inconsistent transcriptional responses to these quite different stimuli, such that the response is more highly correlated between samples than within replicates. With the exception of tissue-specific expression, most samples are more highly correlated with a different condition than they are with other replicates of the same condition. **B**) Principle component analysis. The top 3 most informative principle components describe approximately 50% of the variation, separate the tissue-specific data well, but are unable to clearly distinguish other conditions from one another.

Additional file 13: Table S6 - Pairwise comparison of differentially expressed genes

**Additional file 14: Figure S8.**
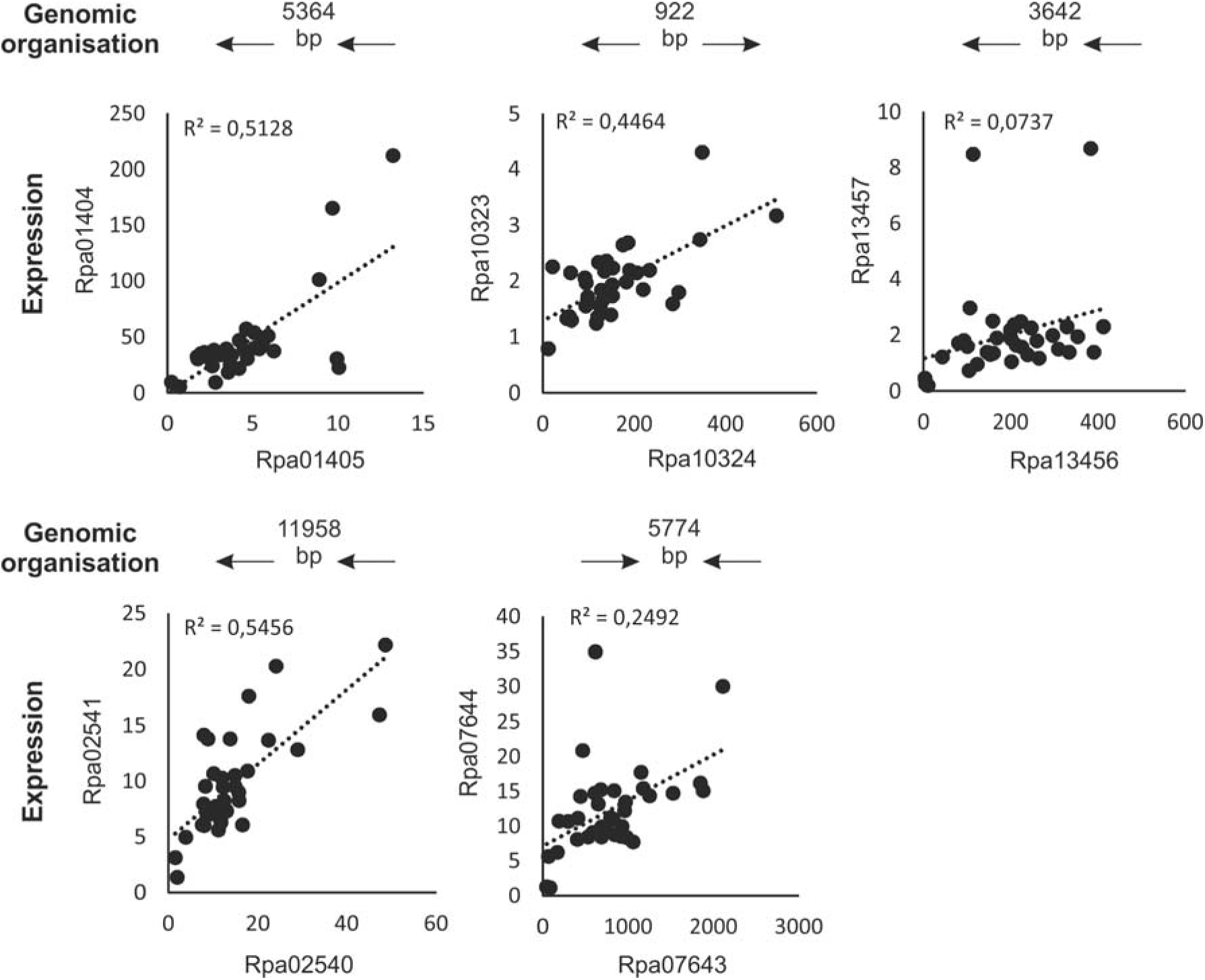
Genomic organisation and expression correlation of all other conserved adjacent pairs in the *R. padi* genome. Five other effectors are present adjacent pairs in all aphids tested. The genomic organisation (arrows), intergenic distance (bp) and expression correlation (R2) are shown for the *R. padi* homologue of each pair.

Additional file 15: Table S7 - Putative contaminants

Additional file 16: File_S1_Alignment_of_123_Single_copy_BUSCO_genes.fasta

## Acknowledgements

We thank Brian Fenton and Gaynor Malloch for kindly providing the *R. padi* and *M. persicae* aphids and for support in generating a *M. cerasi* stock culture. Dr Matt Clark (Earlham Institute) for conceptual design of the sequencing approach.

## Funding

The authors acknowledge the following funding sources: Biotechnology and Biological Sciences Research Council (BB/M014207/1 awarded to SEvdA), European Research Council (APHIDHOST-310190 awarded to JIBB), Royal Society of Edinburgh (fellowship awarded to JIBB).

## Availability of data and materials

Assembled genomes and gene calls are available http://bipaa.genouest.org/is/aphidbase/ and http://mealybug.org. The raw genomic reads for *M. cerasi* and *R. padi* are available at study accession: PRJEB24287 and PRJEB24204, respectively. The raw RNAseq reads for *R. padi* and *M. persicae* reared on host, nonhost and artificial diet at time points 3 hours and 24 hours are available at study accession: PRJEB24317. RNAseq data for *M. cerasi* used to assist gene call is available at study accession PRJEB24338 (paper in preparation). Most custom python scripts used to analyse the data use Biopython, all scripts and methods used throughout this study are available at: https://github.com/peterthorpe5/Methods_M.cerasi_R.padi_genome_assembly https://github.com/sebastianevda/SEvdA_Gephi_array_to_gefx

## Author contributions

PT carried out aphid culture maintenance, collected and prepared material for DNAseq analysis, genome assemblies, annotation, transposons prediction, effector prediction, RNAseq analysis and comparative genomic analysis. CMEM carried out aphid culture maintenance, collected and prepared material for RNAseq analysis. PT and SEVDA analysed the data. PT, SEVDA, and JIBB wrote the manuscript. JIBB conceived and coordinated the project, secured the funding, and contributed to the analyses of data. PJAC assisted and advised on python scripts and conceptual design. DL advised on genome assembly and contamination removal. All authors read and approved the final manuscript.

## Competing interests

The authors declare that they have no competing interests.

## Ethics approval and consent to participate

Not applicable.

